# Time-series metagenomics reveals changing protistan ecology of a temperate dimictic lake

**DOI:** 10.1101/2024.02.09.579653

**Authors:** Arianna I. Krinos, Robert M. Bowers, Robin R. Rohwer, Katherine D. McMahon, Tanja Woyke, Frederik Schulz

## Abstract

**Background:** Protists, single-celled eukaryotic organisms, are critical to food web ecology, contributing to primary productivity and connecting small bacteria and archaea to higher trophic levels. Lake Mendota is a large, eutrophic natural lake that is a Long-Term Ecological Research site and among the world’s best-studied freshwater systems. Metagenomic samples have been collected and shotgun sequenced from Lake Mendota for the last twenty years. Here, we analyze this comprehensive time series to infer changes to the structure and function of the protistan community, and to hypothesize about their interactions with bacteria.

**Results:** Based on small subunit rRNA genes extracted from the metagenomes and metagenome-assembled genomes of microeukaryotes, we identify shifts in the eukaryotic phytoplankton community over time, which we predict to be a consequence of reduced zooplankton grazing pressures after the invasion of a invasive predator (the spiny water flea) to the lake. The metagenomic data also reveal the presence of the spiny water flea and the zebra mussel, a second invasive species to Lake Mendota, prior to their visual identification during routine monitoring. Further, we use species co-occurrence and co-abundance analysis to connect the protistan community with bacterial taxa. Correlation analysis suggests that protists and bacteria may interact or respond similarly to environmental conditions. Cryptophytes declined in the second decade of the timeseries, while many alveolate groups (e.g. ciliates and dinoflagellates) and diatoms increased in abundance, changes that have implications for food web efficiency in Lake Mendota.

**Conclusions:** We demonstrate that metagenomic sequence-based community analysis can complement existing efforts to monitor protists in Lake Mendota based on microscopy-based count surveys. We observed patterns of seasonal abundance in microeukaryotes in Lake Mendota that corroborated expectations from other systems, including high abundance of cryptophytes in winter and diatoms in fall and spring, but with much higher resolution than previous surveys. Our study identified long-term changes in the abundance of eukaryotic microbes, and provided context for the known establishment of an invasive species that catalyzes a trophic cascade involving protists. Our findings are important for decoding potential long-term consequences of human interventions, including invasive species introduction.

## Background

Protists are ubiquitous unicellular eukaryotic microbes capable of widespread dispersal and found globally across most ecosystems, from diverse terrestrial soil environments [1] to all reaches of the ocean [2]. Protists also encompass diverse taxonomic and physiological diversity [3]. Lake ecosystems are no exception, where protists are key players in the microbial loop and determinants of lake trophic status [4]. Like bacteria, protists are essential in the movement of nutrients and energy through an ecosystem [5, 6, 7] but are behaviorally and morphologically more complex [5]. They can display diverse trophic strategies (e.g. mixotrophy [8]) that can alter freshwater food webs [9]. Protists may also colonize macroalgae and assist in carbon transformation, vitamin B12 synthesis, and even silica cycling and storage, functions which have recently been predicted metagenomically [10]. Despite the importance of protists in ecosystem functioning, much less effort has been placed on understanding their biogeography, temporal patterns, and ecology as compared to bacteria. This has been in part due to the visible importance of cyanobacterial blooms [11], and in part due to challenges of defining and differentiating protistan morphology and function [12] or deciphering their complex genomes [5, 12], leaving a large knowledge gap on their geographic extent and lifestyle [13, 5].

Lake Mendota (Dane County, Madison, WI), a temperate dimictic lake, is considered among the world’s best-studied freshwater systems due to the longevity and intensiveness of its monitoring program [11]. Previous studies of Lake Mendota have investigated the seasonal dynamics of the bacterial community in this and nearby lakes [14, 15, 16], including using amplicon sequencing approaches [15] and this metagenomic timeseries [17], but eukaryotic algae and their trophic linkages to bacteria were less frequently considered, though visually identified and counted in microscopy-based surveys [18]. Despite evidence for strong network relationships between bacteria and eukaryotes in both lacustrine and oceanic ecosystems [19], heterotrophic protists are frequently excluded from long-term biomonitoring programs [20], including those in Lake Mendota. Metagenomic sequencing data have the potential to expand the known diversity of lake protists [13], and a recent study showed that 18S rRNA gene data from metagenomes can accurately recapitulate diatom abundance data from sediments [21].

Two invasive species reached high abundances in Lake Mendota during the period from 2000 to 2020 considered in this study. The spiny water flea (*Bythotrephes longimanus*) was first detected in Lake Mendota in 2009 [22], but was shown via sediment cores to have been present more than ten years prior to net tow identification [23, 24]. Zebra mussels (*Dreissena polymorpha*) were detected in the lake in 2015 [25]. Zebra mussels consume phytoplankton, which can result in “benthification” due to enhanced nutrient and light availability in normally shaded benthic lake habitats [25]. New zooplankton predators in lake ecosystems can shunt nutrients from the surface to depth [26]. Trophic cascades can emerge when introduced invasive species eliminate a trophic level, allowing a different taxon to become overabundant [15]. Ongoing microscopic count observations in Lake Mendota confirmed an increase in diatoms (class *Bacillariophyta*) and chlorophytes (phylum *Chlorophyta*) and a decrease in cryptophytes (phylum *Cryptophyta*), but lacked taxonomic resolution or the inclusion of heterotrophic protists that cannot be easily captured by net tows or visually identified [27].

To explore protistan ecology and interactions in Lake Mendota, we analyzed the diversity, structure, and composition of the lake’s protistan community over the course of 20 years. We have extracted 16S and 18S rRNA gene sequences and quantified the abundance of these sequences in the underlying raw read data. We enhanced the taxonomic resolution of Lake Mendota’s eukaryotic microbial community and explored how it has changed in association with prokaryotic community members over a 20-year time series. Further, we introduce the first curated eukaryotic metagenome-assembled genomes (MAGs) from the same long-term metagenomic time series. We show that some of these MAGs can be linked to 18S rRNA gene data, and compare relative abundance of highly complete MAGs in the time series compares to using the 18S rRNA gene. Using enriched metagenomes (metagenomes sequenced from samples with exceptionally high concentrations of the target taxon) sequenced from samples with high abundance of two key metazoans in the lake, *Daphnia pulicaria* (water flea) and *Bythotrephes longimanus* (spiny water flea), we binned MAGs to track environmental abundances of these key metazoans. We tested whether *Bythotrephes longimanus* (the spiny water flea) was present in low abundance in the lake prior to its visual identification in 2009. Our long-term metagenomic study of lake protists demonstrates the power of time series metagenomics to transform our knowledge of freshwater protists and their contribution to lake biodiversity and ecology.

## Methods

### 20 years of metagenome data from Lake Mendota

Lake Mendota is a eutrophic, temperate lake that is heavily influenced by both urban and agricultural land use. The North Temperature Lakes Microbial Observatory (NTL-MO) has collected a time series of water samples starting in 2000 to study microbial communities in Lake Mendota (N 43*^o^*06, W 89*^o^*24). The NTL-MO is coupled with the Long-Term Ecological Research site at Lake Mendota, which collects core limnological data such as water temperature, water chemistry, and plankton microscopy counts (all data available at http://lter.limnology.wisc.edu). NTL-MO collected depth-integrated samples from the epilimnion (approximately 0-12 meters depth), primarily during the ice-free period (approximately March-December) and filtered at 0.2*µ*m prior to storage at -80*^o^*C until further processing.

At the Joint Genome Institute (JGI), 323 of these samples collected from 2000 to 2018 were sequenced on the Illumina HiSeq 2500 platform. The workflow used to process these reads is described in JGI’s metagenomic assembly workflow [28], and includes standard deduplication, filtering, and error correction prior to metagenomic assembly with `metaSPAdes` versions 3.13.0 (*n* = 78) and 3.14.1 (*n* = 245) [29]. The resulting 323 assembled metagenomes across the 20-year time series were processed and annotated using the Integrated Microbial Genomes and Microbiomes (IMG/M) processing pipeline [30]. Additional description of the dataset and processing workflow can be found in [17].

### rRNA gene discovery workflow

To identify small subunit (SSU) rRNA genes in the metagenomic contigs, we used `cmsearch` from the Infernal package (1.1.4) [31] on each assembly with the eukaryotic (RF01960) and bacterial (RF00177) covariance models from the Rfam database [32, 33]. The longest match per contig from each model was retained, and the 16S and 18S rRNA gene sequences were extracted. Identified SSU rRNA gene sequences were filtered by sequence length and clustered at 97% sequence similarity using `VSEARCH` [34]. An absolute minimum cutoff of 500 bp was used for 16S rRNA gene sequences, while a cutoff of 750 bp was used for 18S rRNA gene sequences. We filtered the sequence matches using the National Center for Biotechnology Information (NCBI)’s `BLAST` alignment tool [35] against an alignment database of the rRNA gene sequences from the RefSeq database available from the NCBI’s ftp server [35, 36, 37]. An e-value cutoff of 1e-4 was used, and a minimum alignment length of 100 bp was applied. After obtaining the initial `blastn` hit to the RefSeq rRNA gene database, we used the Entrez Direct command-line utilities to query relevant identifiers from the NCBI database [38, 39]. In order to minimize any issues inherent to rRNA discovery within assembled genomes, namely the fact that rRNA genes may not assemble consistently, in particular for less abundant organisms, we retained only clusters of 16S or 18S rRNA gene sequences which appeared in ten or more datasets out of the total of 323 selected assembled metagenomes for the purposes of the time series trends. Only within-contig matches to the PR2/Silva database that also aligned at the domain level to the appropriate rRNA model (e.g., domain bacteria with RF00177 as the underlying model) were included, and contigs with a qualifying match to both domains were entirely excluded. This condition occurred three or fewer times per assembly, and was likely the consequence of a poor quality sequence to begin with, as these alignments tended also not to meet the alignment length or percentage identity cutoffs of 200 bp and 80%, respectively.

Subsequently, we constructed an 18S rRNA gene and 16S rRNA gene phylogenetic tree separately for eukaryotes and bacteria, respectively using randomly selected sequences from the combined curated database clustered at 85% sequence similarity to approximate order-level groupings [40] as well as the clustered operational taxonomic units (OTUs) from the rRNA gene discovery workflow. All sequences were aligned to the corresponding eukaryotic (RF01960) and bacterial (RF00177) [32, 33] covariance model. A combined alignment file was generated using the `cmalign` tool from the `Infernal` package [31] (1.1.4). `IQ-TREE` (2.1.4) was used to construct a phylogenetic tree using the combined alignment file [41], which was then visualized using `GGTree` in `R` (v. 4.1) [42, 43]. In the case of disagreement in the taxonomic annotation between the cluster representative (the longest sequence in the 97% sequence identity OTU grouping) and the consensus annotation of the cluster members, the placement in the tree was used to determine the more likely annotation of the sequence.

Once we had found a set of contigs containing an SSU rRNA gene, we quantified the abundance of these contigs in the raw sequence files, in order to track the relative abundance of taxa in the dataset over time. We aligned filtered raw reads from the JGI assembly pipeline to the subset of the assembled contigs that had a 16S or 18S rRNA gene recovered. We did this such that all samples were quantified against the same assembled contig set; however it does result in much higher estimated abundance values than when the respective assembly for each set of raw reads is used. We compared this approach to quantification using only clustered rRNA genes and found a *>* 66.9% R^2^ (*p <<* 0.01) correlation coefficient between the two, with much of the difference being driven by the fact that many samples did not contain the 16S or 18S rRNA gene when only clustered genes extracted by each assembly were used. Alignment was done using the `Bowtie 2` aligner (2.4.4) [44], then BAM, sorted BAM, and SAM files were generated using `SAMtools` [6] (1.3) and duplicated reads were marked and removed using `Picard` [45] (2.26.9). Finally, `SAMtools` [6] (1.3) was used to obtain coverage estimates and a number of mapped raw reads per contig. This final number of mapped reads per contig was used with the contig length to calculate a per-contig version of transcripts per million (TPM) [46], which has recently been adopted for metagenomic data [47]. We call this version of TPM “CPMcontig”. We calculated ‘CPMcontig using the following formula:

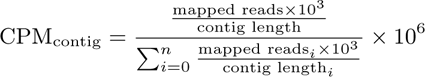

where *n* is the total number of contigs in the assembly. The total sum of CPMcontig in a given assembly will hence always be equal to 10^6^.

Because of inconsistencies in 16S and 18S rRNA gene copy number between different taxa (e.g., [48, 49, 50]), we further normalized abundance estimates to Z-scores during the period of interest. While not making disparate samples directly comparable [46, 51, 52], Z-scores standardize measurements already subject to variance stabilization via CPMcontig calculation and simplifies interpretation of changes in abundance during the sampling period. Instead of using raw abundance, the Z-score of each abundance estimate was calculated as a relative measure of the change in the size of the community of the respective taxonomic group over time. We also took into account unbalanced sampling over the total sampling period by using the weighted mean and weighted standard deviation within climatological seasons (the ice period, spring, early summer, late summer, and fall) as defined by Rohwer et al. (2022; [15]). We took unbalanced sampling into account by dividing by the total number of samples observed during each season and year to avoid unduly biasing overall abundance estimates towards years that were sampled more densely.

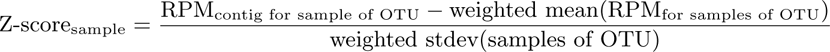

The Z-score metric constrains the range of values for multiple species over the twenty years of measurements [53] and partially addresses the limitations of TPM (thus, RPMcontig) for between-sample comparisons [46, 52].

We conducted weighted two-sample t-tests (using the R package weights version 1.0.4 [54]) with a Benjamini-Hotchberg correction with a 0.05 *alpha* value as separated between the pre-invasion of spiny water flea (2009 and before) [22, 23] and post-invasion (after 2010) segments of the time series and their corresponding abundance estimates in genes per million to categorize each eukaryotic clustered 18S rRNA gene.

### Identifying co-abundant community members in time

In order to identify frequently co-occurring organisms on the basis of the abundance of their 18S and 16S rRNA gene sequences over the full twenty year time series, we used hierarchical clustering. We clustered the sequences with the `hclust` function from the `stats` package within `R`, and created a cluster dendrogram using the Ward method and the Canberra distance function [43]. We used the `pvclust` function for uncertainty estimation and bootstrapping with 1000 bootstrap iterations [55], and an initial adjusted approximately unbiased bootstrap probability value (AU) of 0.80 for preliminary filtering of cluster results. Visualization was performed using the `dendextend` package [56].

We also conducted a broader co-abundance network analysis between all taxa using all-by-all Pearson correlations as computed using the `rcorr` function within package `Hmisc` [57]. To control for multiple testing and identify significant Pearson correlations between taxa, we used the Benjamini-Hochberg procedure with a false discovery rate of 0.25 and an alpha value of 0.05 [58]. In our network visualizations, we use a correlation coefficient cutoff of 0.5 to identify correlations that are both significant and strong. Network visualization was performed using the `network` and `igraph` packages within R version 4.1.0 [43, 59, 60]. The Louvain clustering algorithm as implemented in `igraph` was used to identify coherence communities within the network [61, 60].

### Eukaryotic metagenome-assembled genomes (MAGs) from Mendota metagenomes highlight key groups of protists

#### Eukaryotic metagenome-assembled genome identification, taxonomic annotation and phylogeny

Each individual assembled metagenome from discrete sampling dates was scanned for eukaryotic sequences with `EukRep` [62] (0.6.7). Identified eukaryotic sequences were combined into bins using `METABAT2` [63] (2.12.1) and gene prediction was performed using `Prodigal` (2.6.3) [64]. Completeness and contamination of MAGs was estimated using `EukCC` [65] (0.2).

We used the BUSCO (Benchmarking Universal Single-Copy Orthologs, [66]) genes found in 60% of MAGs of completeness *>* 40% as well as those same genes from a selection of reference genomes and transcriptomes (73 reference genomes sequenced and curated by the JGI and 35 eukaryotic transcriptomes from the Marine Microbial Eukaryote Transcriptome Sequencing Project (MMETSP; [67]) to align the highly complete MAGs and construct a phylogenetic tree. The highly complete MAGs each had more than 90% of the genes found in 60% of all highly complete MAGs.

The above workflow only retained four of the identified MAGs, so we constructed a secondary phylogenetic tree using MAGs with completeness *2:* 10% that had 30% or more of the BUSCO genes that were found in 15% or more of the MAGs. This tree was constructed using this subset of BUSCO genes found with differing consistency *2:* 30% among the MAGs.

In addition to extracting MAGs from the time series metagenomes, we also did so for two enriched metagenomes, one enriched with *Daphnia pulicaria* and one enriched with *Bythotrephes longimanus*, the spiny water flea. Enrichment was performed by physically separating each zooplankton group from a water sample and sequencing zooplankton-dominated material. The raw read sequencing data for these two enriched metagenomes is available online at the NCBI’s Sequence Read Archive (SRA) under accession number *to be added*. We used the same workflow for these MAGs, including completeness estimation with `EukCC` as described in the Methods.

Using the extracted BUSCO proteins for each of the MAGs and a number of reference genomes curated by the JGI [68], we used trimal [69] to reduce spurious sequences, Clustal Omega [70] (1.2.4) to perform multiple sequence alignment, and FastTree (2.1.10) and RAxML (8.2.12) [71] for phylogenetic analysis.

For taxonomic annotation of the MAGs, we compared three approaches. First, we used the suggested NCBI database and utility provided by EukCC ([65]; 0.2) in concert with completeness estimates Second, we used `EUKulele` [72] with a custom database comprised of the MMETSP [67, 73], MarRef [74], and a selection of reference genomes of eukaryotes from the JGI’s Genome Portal [68] to annotate the proteins extracted from the contigs binned into each MAG.

#### Eukaryotic MAG abundance quantification

The abundance of the top 51 most complete MAGs was quantified using the output of the same workflow as for the rRNA gene sequences to which the filtered raw reads from the JGI `metaSPAdes` [29, 75] assemblies were mapped. In brief, raw reads were aligned to contigs from full metagenomic assembly using the `Bowtie 2` (2.4.4) [44], then BAM, sorted BAM, and SAM files were generated using `SAMtools` [6] (1.3) with duplicates marked and removed using `Picard` [45] (2.26.9). Contigs from the MAGs were searched against the full metagenomic assembly to identify MAG representatives in each sample using `mmseqs2 search` with a sensitivity parameter of 0.70 [76]. Matches were filtered for those with e-value ≤ 10*^-^*^50^ and percentage identity of 0.90 or above. The sum of the abundance of those clustered contigs was taken for each sample to be the per-sample MAG abundance. `SAMtools` (1.3) was used to obtain coverage estimates and a number of mapped raw reads per contig [6]. This procedure was used in place of mapping the MAGs directly to ensure that an appropriate value for CPMcontig could be established using the full set of contigs extracted from each sample. This avoided bias related to the sample that each MAG was originally extracted from (since all MAGs were treated using the same procedure) and the total number of MAGs tested. Bias related to the total size of each MAG was unavoidable. Using a sequence search between binned contigs and all the contigs minimizes bias because quantification is done only based on assembled sequences, but bins with larger numbers of contigs will inevitably have more opportunities for a sequence match to be made that meets the selection criteria.

### Phylogenetic analysis for individual clades

Phylogenetic analysis for individual clades was performed similarly to the general phylogenetic analysis for eukaryotes and bacteria. In brief, using clustering with `VSEARCH` [34], alignment with Infernal [31], and phylogenetic analysis using `IQ-TREE` [41] (2.1.4) and plotting in `ggTree` [42].

### Putative bacterial symbionts of microeukaryotes

Several bacterial taxa are known endo- or ectosymbionts of eukaryotic microorganisms. We identified representatives of these taxa from the bootstrapped hierarchical clustering procedure described in the “sec:cooccurrence” subsection. In order to explore the bacterial taxa which fell into clusters with defining temporal trends, we calculated the correlation between the Z-scores (as calculated over the full time series) of the putative endosymbiont clades and all eukaryotic OTUs, selecting OTUs with Pearson correlation coefficients of at least 0.5 as explained by the co-abundance network analysis, and which co-occurred in at least 10 distinct samples with the putative endosymbiont OTU. We also explored the number of samples in common between each bacterial taxon and co-clustering eukaryotic taxa, identifying the proportion of the time that these bacterial taxa were present independently of their eukaryotic counterparts.

## Results

### Microbial communities in Lake Mendota show seasonal partitioning

A total of 606 clusters of 18S rRNA gene sequences and 2,223 clusters of 16S rRNA gene sequences at 97% sequence similarity were recovered from the 323 assembled metagenomes. These were then further filtered by those with ten or more occurrences in the dataset and high quality `blastn` matches to a total of 109 18S rRNA gene sequences (Figure 1) and 608 16S rRNA gene sequences (Supplementary Figure 3). The average length of filtered 18S rRNA gene sequences was 1,834 *±* 363 bp, while the average length of 16S sequences was 1298 *±* 299 bp. The eukaryotic OTUs spanned major branches of eukaryotic life, including *Metazoa* (including invertebrate arthropods like *Daphnia* and *Bythotrephes longimanus*, the water flea and spiny water flea, respectively), *Apicomplexa* (parasites), *Cryptophyceae* (cryptophyte algae), and *Ochrophyta* (a group that includes diatoms - class *Bacillariophyta* - and other algae). Some of these sequences, including many cryptophytes and arthropods, were independently assembled and recovered from the majority of the Lake Mendota metagenomes (Figure 1). As expected based on the Plankton Ecology Group model [77, 78], some eukaryotic taxa also showed strong seasonal patterns, such as the tendency of cryptophytes to be most abundant in winter, and the tendency of diatoms (labeled Ochrophyta, class *Bacillariophyta*) such as family *Stephanodiscaceae* to be abundant in fall and spring ([79], Figure 1,2). Most microeukaryotic taxa were most abundant in spring, the ice-on period, and fall (Figure 2A). However, arthropods, including most zooplankton taxa, were most abundant in the clearwater period (mid-spring to early-summer, when phytoplankton abundances decline; Figure 2A), coinciding with the lower number of distinct SSU rRNA genes identified (alpha diversity; Figure 2B and C). Both bacteria and eukaryotes had highest taxonomic diversity in the fall and winter according to SSU rRNA gene recovery (Figure 2B and C). Bacteria had universally higher alpha diversity than eukaryotes, but eukaryotic diversity was highest relative to bacterial diversity in the month of April during the spring bloom (Figure 2B). Copepods had higher relative abundance in winter, rotifers had higher relative abundance in summer and fall, and cladocerans had higher relative abundance in spring and summer (Figure 5).

**Figure 1:**
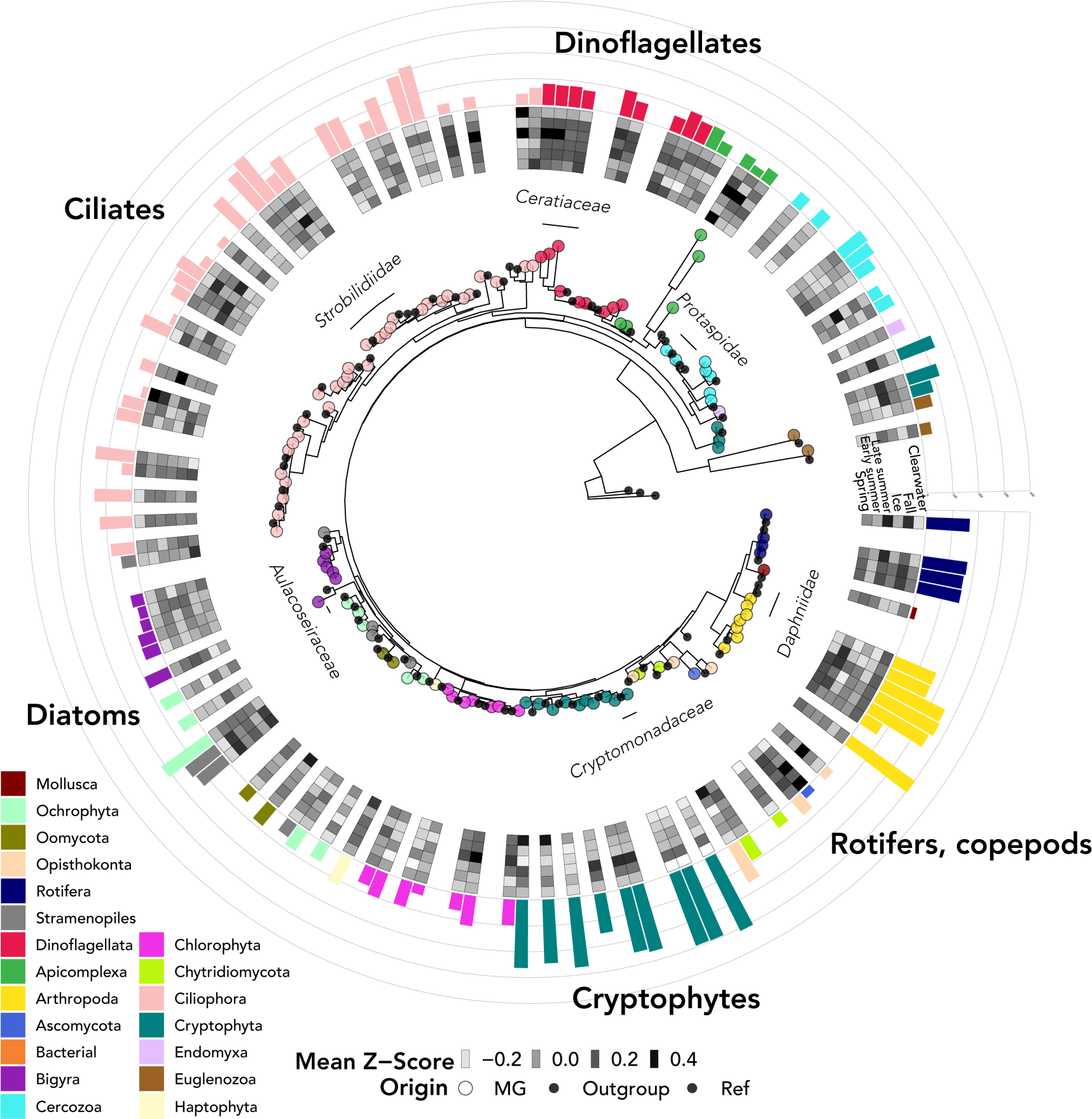
Phylogenetic tree of 18S rRNA gene sequences extracted from Lake Mendota metagenomes over the twenty-year time series. Colored, filled circular points on the tree indicate novel sequences extracted from the Mendota metagenomes, while smaller black circles denote a representative set of previously published sequences. The rings of the heatmap represent the mean Z-score of extracted sequence during each of the seasons of the year, and bars show the number of samples the 18S rRNA gene was found in.

**Figure 2:**
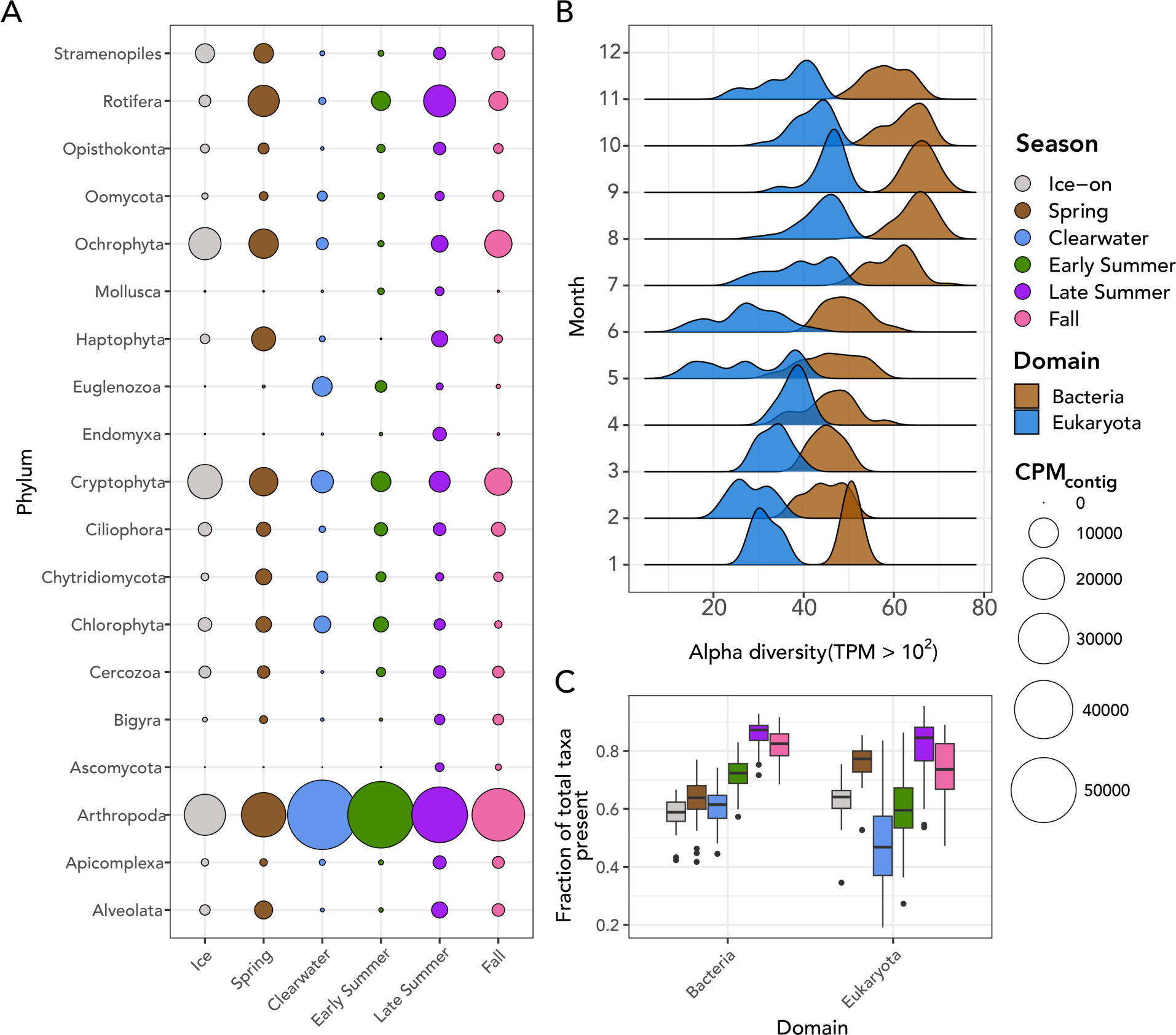
Evidence of seasonal variability of SSU rRNA genes. A: The abundance of major taxa of protists and metazoans in Lake Mendota by season. Circles are colored by the season of the year and sized according to total abundance via CPM_contig_ for that month. B: Alpha diversity as assessed by extracting OTUs with TPM of 100 or greater. Distributions of alpha diversity are shown for both eukaryotes and bacteria for each month of the year in the time series. C: Proportion of total observed taxa present in each season of the dataset for (left) bacteria and (right) eukaryotes.

### Changes in protistan community composition: an increase in *Alveolata*

#### and a decrease in cryptophytes

In the second half of the time series (2010-2019), several taxa showed significant increases in relative abundances (Figure 3). This occurred following the 2009 invasion of *Bythotrephes longimanus* (spiny water flea) [22]. 59 vetted eukaryotic OTUs did not show an increase in abundance between the periods of before 2010 and after 2010, while 42 OTUs increased, including 28 alveolates, 7 dinoflagellates of class *Dinophyceae*, 14 ciliates of class *Ciliophora*, and 3 candidate Apicomplexan parasites (Figures 3 and 4). A significant increase in several metazoan OTUs was observed in the dataset, including an OTU assigned to the Zebra mussel (genus *Dreissena*; *p* = 8.2*e^-^*^4^), which tended to be low or zero abundance for the majority of the time series but increased markedly following its documented irruption in Lake Mendota (2015; Figure 5; [25]), four copepod OTUs of order *Cyclopoida* (2 with *p <* 2*e^-^*^5^, 1 with *p <* 3*e^-^*^3^, 1 with *p ⇡* 0.01), a rotifer of order *Flosculariaceae* (*p <* 2*e^-^*^5^), and two rotifers of order *Ploima* (*p <* 2*e^-^*^5^) (Figures 3 and 5). Ciliate classes *Nassophorea*, *Litostomatea*, and *Heterotrichea* and dinoflagellate class *Dinophyceae* had statistically significant increases in 2010-2019 relative to 2000-2009 (Figure 4B,D). *Nassophorea*, *Litostomatea*, and *Dinophyceae* also showed higher relative alpha diversity during this period (Figure 4E), meaning that more of the total OTUs belonging to each class were recovered with sufficient abundance in the second half of the time series.

**Figure 3:**
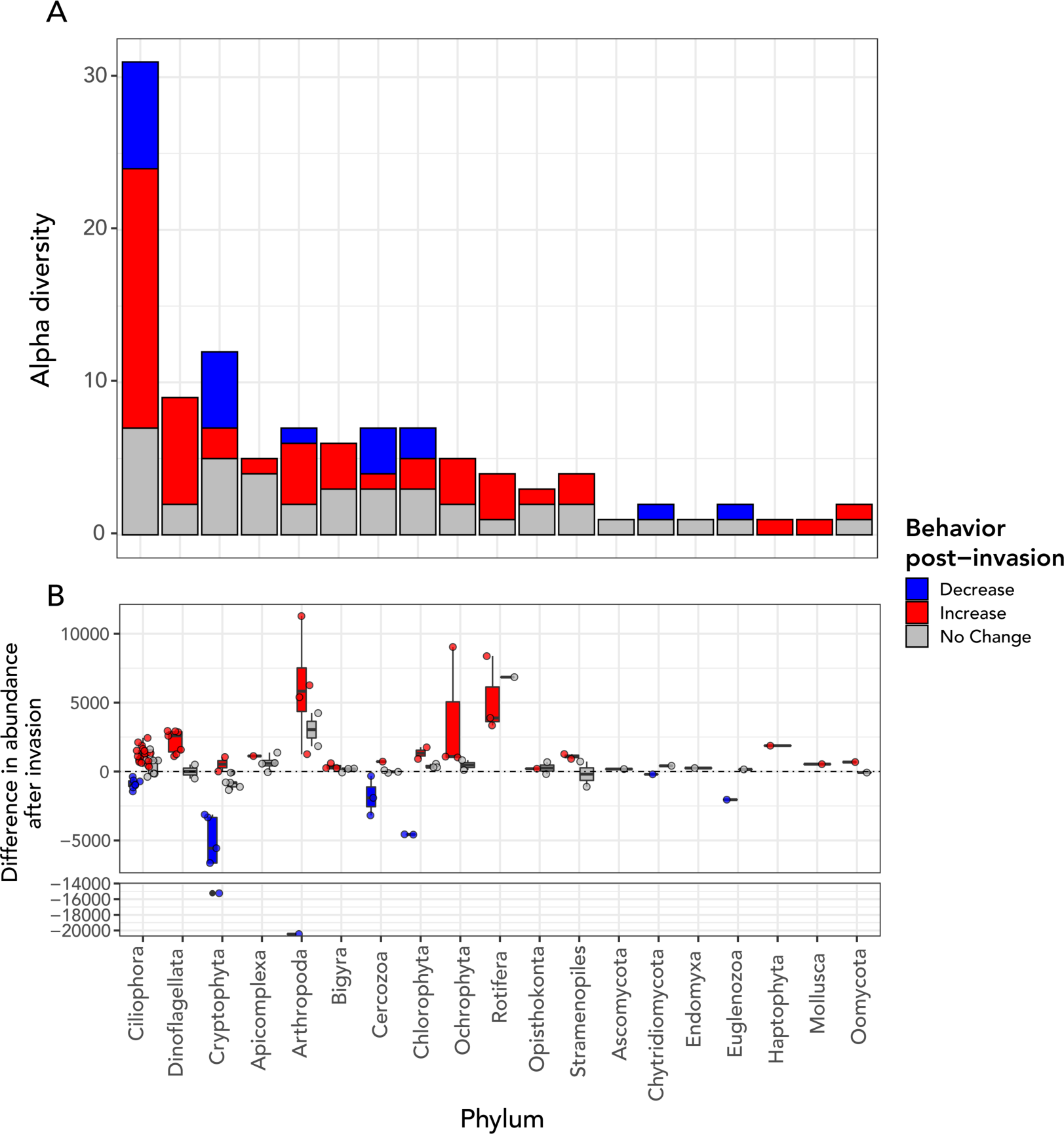
Shift in abundance of microbial eukaryotes and other relevant eukaryotic taxa after the spiny water flea invasion. A: Phylum breakdown of OTUs that did or did not show statistically significant increases in CPM_contig_ before and after 2010. The majority of statistically significant increases were from OTUs annotated as clade Alveolata (including phyla Ciliophora, Apicomplexa, and Dinoflagellata). B: Difference in mean CPM_contig_ after the invasion for OTUs and mean CPM_contig_ before the invasion of the spiny water flea, grouped and colored by the statistical significance of the change in abundance.

**Figure 4:**
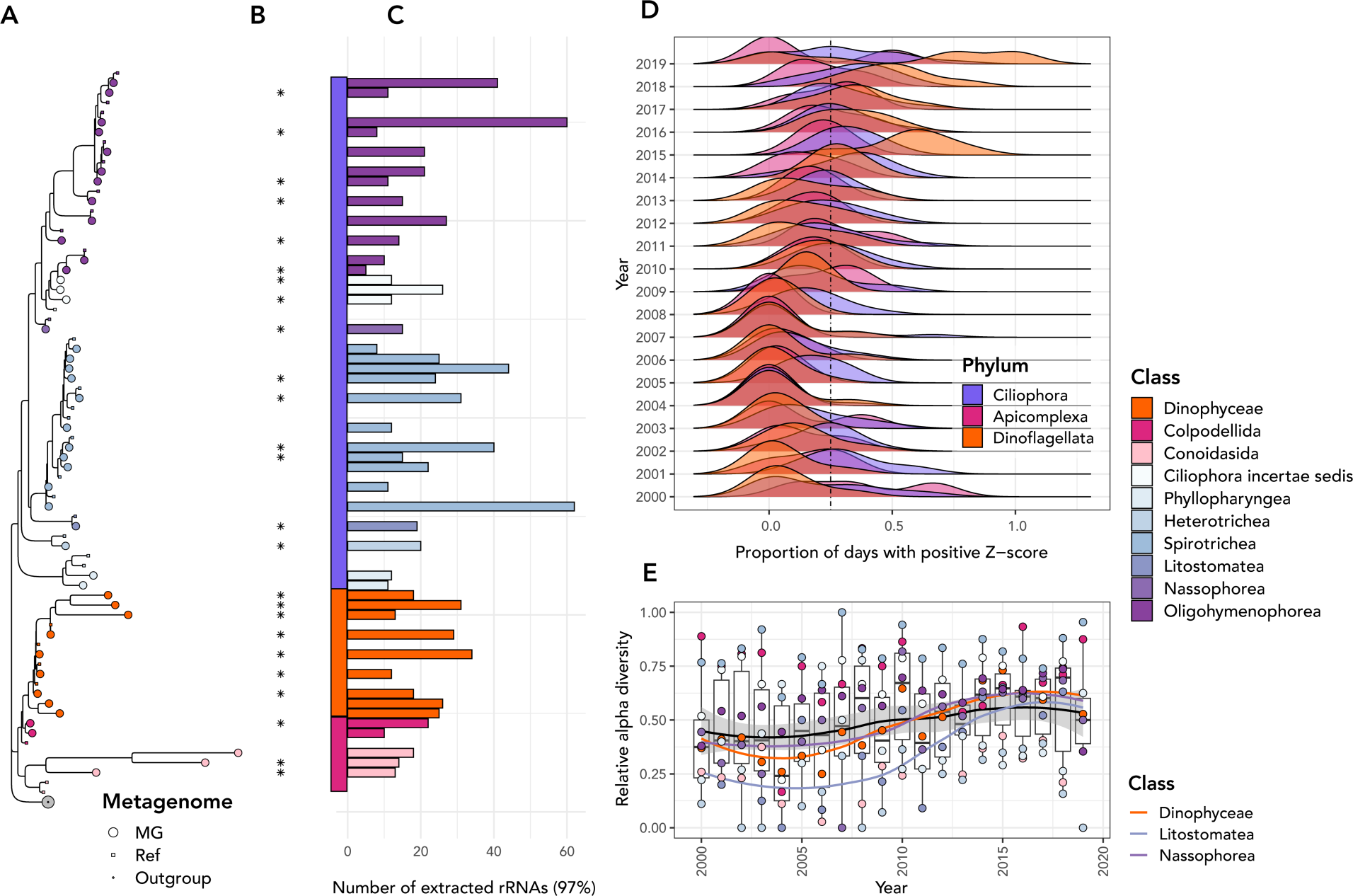
Abundances of Alveolates in the Lake Mendota time series. A: Phylogenetic tree showing all alveolate (clade Alveolata) sequences (square tip points) alongside their NCBI RefSeq closest blast matches (small circles), highlighted by their taxonomic Class. The green circle at the bottom of the tree is an outgroup clade of *Ochrophyta* used as an outgroup. B: Extracted 18S rRNA gene sequences are labeled with a “*” if they changed significantly in abundance according to the procedure described in the text. C: The number of 18S rRNA genes from all samples that fell into the cluster (97% sequence identity) in the phylogenetic tree. Ciliates of class *Spirotrichea* tended to be most highly abundant in the dataset. D: Distribution of days that had positive Z-scores in abundance for each Alveolate phylum for each year of the dataset. In the later years of the dataset, dinoflagellates in particular had a higher proportion of total sampling days that had elevated abundance relative to the mean. E: Alpha diversity (number of OTUs) present relative to total extracted for each alveolate class in each year of the dataset. Classes *Nassophorea*, *Litostomatea*, and *Heterotrichea* were universally more commonly present in the latter half of the dataset (2010-2019), and in particular in 2012-2018. Lines drawn on top of the boxplots and point show the overall trend (black) and colored trends in relative alpha diversity for the three taxonomic classes for which alpha diversity trends were significant.

The remainder of the OTUs that showed significant increases in abundances were two Stramenopiles (order Hyphochytriaceae), two Chlorophytes (order Chlamydomonadales), and a Haptophyte (order Prymnesiales).

By contrast, all eukaryotic taxa that showed statistically significant decreases after 2010 belonged to three phyla: Cryptophyta, Ciliophora, and Cercozoa. This included two unclassified Cercozoans (*p<* 0.005), two ciliates of class *Spirotrichea*, and five cryptophytes. This was 41.7% of all cryptophyte OTUs recovered (12 total), and 100% of the cryptophyte OTUs annotated to belong to taxonomic order *Cryptomonadales* (*n*=2, *p<* 0.001).

### Putative relationships between eukaryotic and bacterial taxa

We leveraged the metagenomic rRNA gene sequences to identify clusters of eukaryotes and prokaryotes in Lake Mendota (Figure 7) and further to explore co-abundances of eukaryotes with prokaryotes, and hence predict their putative relationships. To identify these clusters, we used both hierarchical clustering and network analysis of the rRNA gene sequences. The first set of organisms we focused our analysis on were bacteria known to be potential endosymbionts of protists. We identified six OTUs of bacteria that were affiliated with known endosymbionts of microbial eukaryotes based on their annotated taxonomy (Supplementary Figure 8; complete list of correlations and hierarchical clustering modules available in Supplementary Tables 1 and 2). Two endosymbiotic genera showed statistically significant (*p<* 0.01) increases in abundance after the spiny water flea invasion: a *Caedimonas* OTU from order *Holosporales* and a *Rickettsia* OTU from order *Rickettsiales* [80]. Importantly, these bacterial OTUs were significantly correlated with eukaryotic OTUs via hierarchical clustering and/or network analysis (Figure 7). Network analysis revealed that many ciliate (phylum Ciliophora) OTUs had multiple correlations to proteobacterial OTUs, including two families that contain known endosymbionts: a *Rickettsiales* (r=0.57) OTU and a *Holosporales* (r=0.60) OTU. Two ciliates formed a module with a *Sphingobacteriaceae* bacterium (*Bacteroidetes*; AU *p*-value *<*0.01). One diatom (phylum *Ochrophyta*; class *Bacillariophyta*) OTU was significantly correlated with an OTU belonging to the family *Anaplasmataceae* (order *Rickettsiales*; r=0.72), We also identified bacterial taxa that were more likely associated with eukaryotes as part of local environmental conditions. One eukaryotic OTU annotated as *Chytridiomycota* was significantly correlated with a *Planctomycetes* (*Isosphaerales*) OTU (r=0.66). A diatom OTU was significantly correlated with an OTU of cyanobacterium *Chamaesiphonaceae* (*Synechococcales*; r=0.87), and a cyanobacterial OTU of *Microcystaceae* (*Chroococcales*; r=55). A dinoflagellate (*Peridiniales*) was correlated with a bacterium of order *Saprospirales* (r=0.54) and another bacterium of order *Sphingomonadales* (r=67). One cluster of ciliate *Nassulida* contained bacteria of orders *Desulfuromonadales*, *Synechococcales*, and *Phycisphaerales* (AU *p*-value*<*0.01). Other relationships may have been associated with direct trophic relationships, for example the correlation between a dinoflagellate (*Peridiniales*) with a cercozoan (*Cryomonadida*; *p*¡0.001, r=0.58) and a ciliate (*Oligotrichia*; *p*¡0.001, r=0.60). Further, we identified bacterial OTUs known to be potential constituents of protistan microbiota or members of biofilms [81, 82]. These included an OTU of *Borreliaceae* within order *Spirochaetales* which significantly correlated to a diatom (r=0.75), and one of cyanobacterium *Chamaesiphonaceae* (*Synechococcales*), also correlated to a diatom with the strongest correlation coefficient value recorded in the dataset (r=0.87; Figure 10; Supplementary Figure 7). Chlorophytes (green algae) were correlated with two *Comamonadaceae* (*Burkholderiales*) OTUs (r=0.50 and r=0.51), a *Rhodobacteraceae* (*Rhodobacterales*) OTU (r=0.53), a *Chthoniobacteraceae* (*Chthoniobacterales*) OTU (r=0.52,0.53), a *Flavobacteriaceae* (*Flavobacteriales*) OTU (r=0.51,0.52), and a *Sphingosinicellaceae* (*Sphingomonadales*) OTU (r=54) (Figure 10). Some of these bacterial OTUs were also contained in a large hierarchical clustering module which contained several eukaryotes: two chlorophytes, a chrysophyte, a dinoflagellate, an apicomplexan, three ciliates, and two cercozoans (Figure 10; Supplementary Table 2). Nine bacteria were contained in the same module as this cluster of eukaryotes, including two alphaproteobacteria (*Sphingomonadales* and *Rhodospirillales*), and two planctomycetes (Supplementary Table 2).

### Eukaryotic MAGs match several abundant taxa in 18S rRNA gene time series

A total of 4,511 eukaryotic MAGs were recovered across all samples, of which the majority were small and incomplete, and only 154 were sufficiently large for `EukCC` [65] to calculate a completeness estimate. 14 of these had greater than 20% completeness, 5 had greater than *>*50% completeness, and 2 had above *>*75% completeness. Leveraging `EukCC` completeness estimates, the 10 MAGs that were at least 40% complete had 28 BUSCO genes from which a 90% subset could be found in at least 60% of the 40%-complete MAGs. 10 MAGs met the second threshold criterion described in the methods and were included in the second phylogenetic tree alignment. In practice, the 10 MAGs which met the shared BUSCO criterion had completeness of 17% or greater. A total of 86 BUSCO genes were used for the alignment.

We found a chlorophyte (green algal) MAG which had both an associated 18S rRNA gene and was annotated as a chlorophyte and aligned within the chlorophyte clade in the MAG phylogenetic tree (Figure 6A). While its 18S rRNA was part of a cluster that only appeared 6 times in the dataset and hence was excluded from the time series analysis, it had 96.5% identity and a 1,785-nucleotide alignment length to a chlorophyte of genus *Monomastix* via NCBI RefSeq sequence (accession number AB491653). The vast majority of eukaryotic MAGs did not have a co-binned 18S rRNA gene sequence from the SSU rRNA discovery procedure. The most complete MAGs extracted belonged to phylum Haptophyta (Figure 6) and appeared to likely belong to genus *Chrysochromulina*, a documented haptophyte genus in lake ecosystems [83].

**Figure 5:**
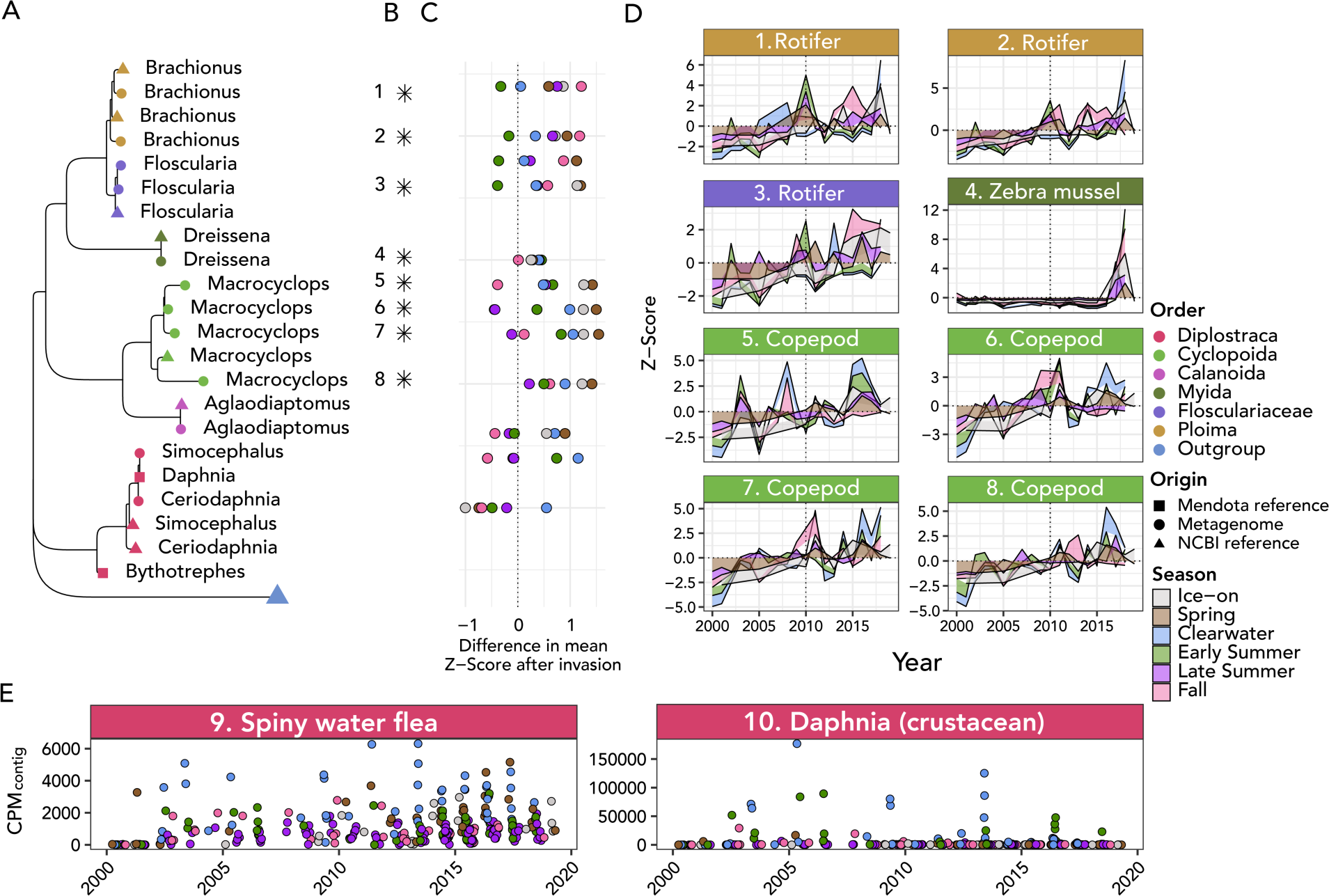
Changing community profile of metazoans in Lake Mendota. A: Phylogenetic tree showing genus-level taxonomic assignment of each metazoan 97% OTU. Shapes indicate sample origin and are colored based on order-level taxonomic assignments of the respective genus. Squares correspond to references extracted from companion metagenomes for *Daphnia* (water flea) and *Bythotrephes* (spiny water flea; references for each generated using enriched metagenomes from Lake Mendota). Tree is rooted at the phylum *Ochrophyta*. B: Stars (“*”) denote significant abundance increases of rRNA clustered OTUs after 2010. C: Difference in average Z-score of abundance of metagenomic rRNA before and after the spiny water flea invasion by year and season. The x-axis corresponds to Z-score, and points are colored by season. D: Time series seasonal Z-scores for taxa numbered in Panel B. E: Mapped abundances of each of the 18S rRNA genes extracted from the companion metagenomes for *Bythotrephes* and *Daphnia*.

#### MAG abundance quantification shows similar patterns to 18S rRNA gene survey

We quantified the abundance of the top 20 MAGs of highest completeness per `EukCC`. Several of these MAGs were annotated as *Haptophyta*, specifically as *Chrysochromulinaceae* (in the most complete Haptophyte MAG, 29.5% of contigs in the MAG showed consensus for *Chrysochromulinaceae*, as opposed to 52.3% for order *Prymnesiales*, 87.5% for class *Prymnesiophyceae*, and 90.2% for phylum *Haptophyta*; MAG IDs ME-TYMEFLIES-MAG 32, 36, 37, 38, 42, 43, 45, and 46; Figure 6). All of these MAGs showed the same pattern of statistically significant increase in abundance in the metagenomic contigs after the spiny water flea invasion (*p <* 0.001). The highest overall abundance was associated with MAGs of highest completeness, though all of the MAGs revealed the same trend of increasing abundance.

Putative chlorophyte MAGs included ME-TYMEFLIES-MAG 33, 47 (70.0% `EUKulele` consensus; 31.4% for class *Mamiellophyceae*; 16.2% for order *Mamiellales*), 48 (86.5% consensus for phylum *Chlorophyta*; 39.4% consensus for class *Trebouxiophyceae*; 26.6% for order *Chlorellales*), and 50 (71.5% `EUKulele` consensus for phylum *Chlorophyta*; 32.6% for class *Mamiellophyceae*). The only putative cryptophyte MAG was ME-TYMEFLIES-MAG 34 (`EUKulele` consensus 79.5% for phylum *Cryptophyta*; 78.5% for class *Cryptophyceae*; 55.7% for order *Cryptomonadales*). Putative ciliate MAGs included ME-TYMEFLIES-MAG 40 (35.5% consensus) and 44 (37.7% consensus).

Putative ochrophyte MAGs included ME-TYMEFLIES-MAG 35 (23.3% consensus), 39 (14.3% consensus), 41 (20.3% consensus), 49 (90.6% consensus for Ochrophyta; 88.3% consensus for class *Bacillariophyta*; 21.6% consensus for order *Thalassionemales*), though it should be noted that all of these MAGs apart from MAG 49 had conflicting annotations between dinoflagellates and ochrophytes, and only MAG 49 was complete enough to be included in the phylogenetic tree constructed using BUSCO genes (Figure 6). In MAG 49, the strong signal of an exceptionally large diatom bloom near the end of the time series was reproduced by the contigs that clustered with the MAG, although the annotation provided by the 18S rRNA gene sequence was order *Aulacoseirales*, which is another taxonomic order of pennate diatom. MAG 39 also showed a substantial, though less than 10% of the CPM maximum of MAG 49, increase in abundance during the same time period (Figure 6).

Further, we used an auxiliary dataset, one of which was enriched in cladoceran *Daphnia pulicaria* and the other in spiny water flea *Bythotrephes longimanus*, as a comparison. We extracted 18S rRNA gene sequences from these datasets and binned MAGs. `EukCC` estimated the completeness of the *Daphnia pulicaria* MAG at 32.4% with 8.8% contamination, and the *Bythotrephes longimanus* MAG at 38.2% completeness and 8.8% estimated contamination (Figure 6).

**Figure 6:**
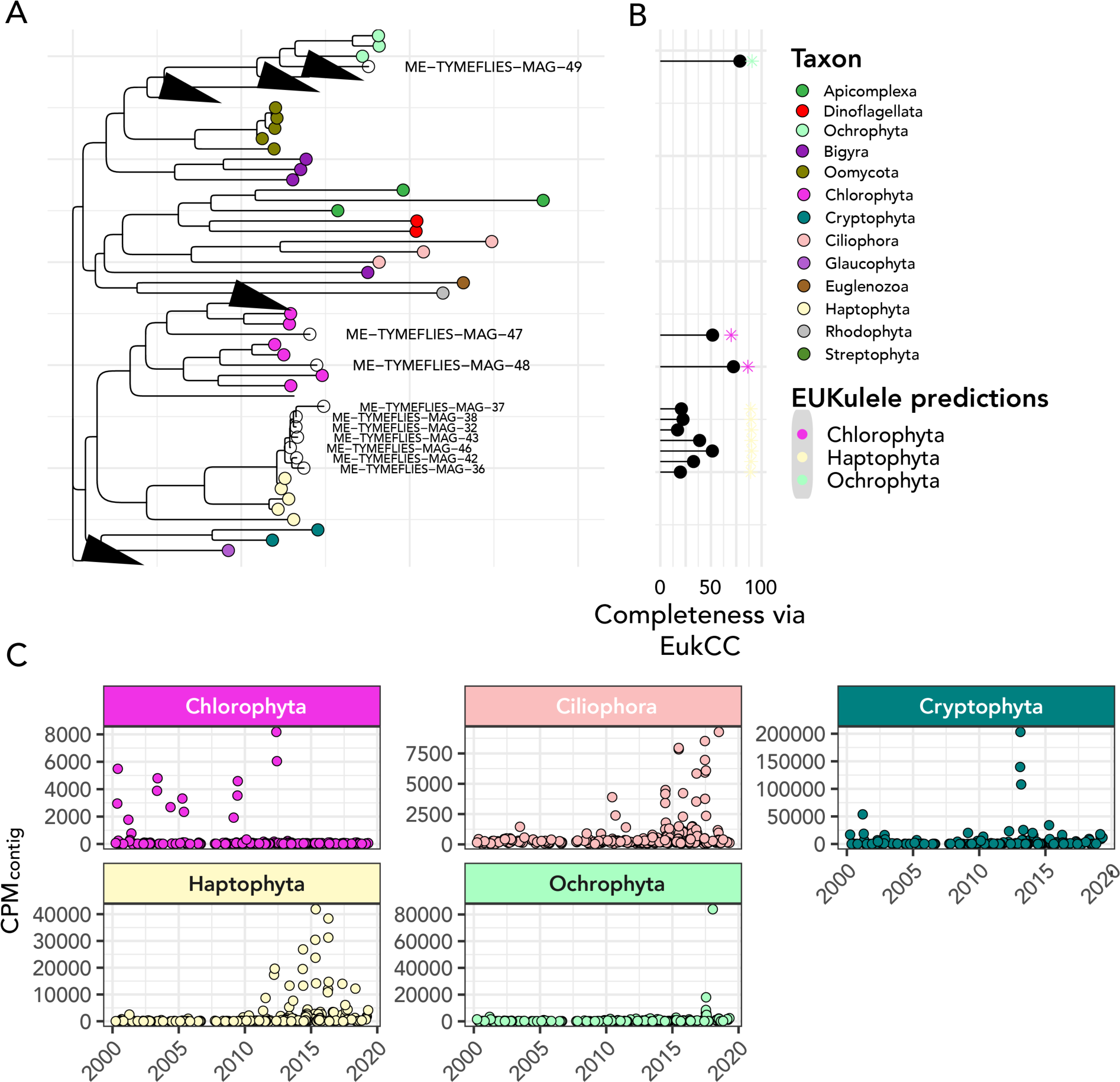
Metagenome-assembled genomes recovered from Lake Mendota span a wide range of eukaryotic taxa. A: Phylogenetic tree of extracted MAGs using concatenated alignment of BUSCO genes alongside reference genomes from the corresponding phyla. Only MAGs which contained a sufficient number of BUSCO genes (see Methods) were included. Unfilled and labeled circles correspond to MAGs extracted from the Lake Mendota time series. B: Lollipop plot showing completeness of each MAG via EukCC (connected point) and the percentage consensus of the phylum-level taxonomic annotation provided by EUKulele (colored asterisk (*)). C: Abundance of the most abundant metagenome-assembled genome extracted from within each taxonomic group across the metagenomic time series, expressed in CPM_contig_.

## Discussion

Our results show that 18S rRNA genes can be recovered from metagenomic time series data and used to interpret temporal patterns. These patterns combined with even coarse taxonomic information provided ecological insights into protists, a diverse and abundant group of organisms frequently ignored by freshwater microbial ecologists. We identified eukaryotic lineages that significantly increased over the course of the time series, and conducted a network analysis to predict potential associations between protists and bacteria. Major changes in abundance were observed in Alveolates (including a selection of dinoflagellates, ciliates, and Apicomplexans; no phytoplankton of clade *Alveolata* are cataloged in the LTER dataset for Lake Mendota), some diatoms, as well as some classes of metazoans (*Opisthokonta*), including genus *Dreissena* (Zebra mussel) and several copepods, and decreases in several OTUs annotated to be cryptophytes. These changes occurred following the invasion of the spiny water flea, *Bythotrephes longimanus*, near the end of 2009, which significantly changed the ecology of Lake Mendota and likely contributed to these shifts in protistan community structure [84]. Increases in eukaryotic phytoplankton may be attributable to a decrease in grazing by *Daphnia* predators due to *Bythotrephes* invasion, as previously established in the literature [23, 22, 84, 85].

**Figure 7:**
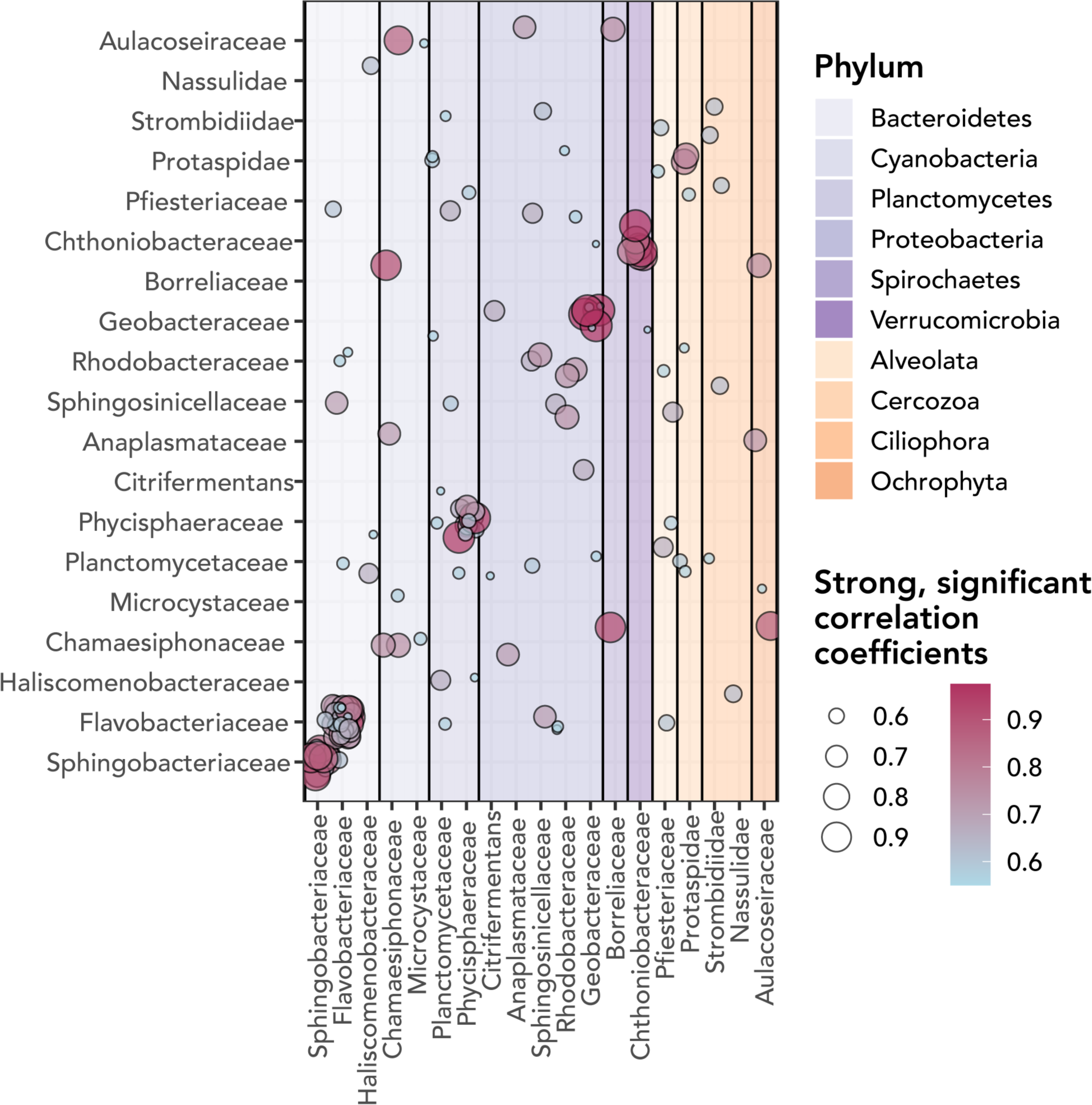
Network analysis over 20-year time series of metagenomes suggests stable connections between eukaryotes and bacteria. Circles correspond to significant, strong correlations between taxa listed on the x- and y-axes, and are both colored and sized according to the correlation coefficient. Only correlations with a significant Benjamini-Hochberg-corrected p-value and a correlation coefficient of greater than or equal to 0.50 are included visually. The shading corresponds to whether the x-axis partner is a bacterial or a eukaryotic phylum. Self-correlations are excluded when the small subunit rRNA gene sequences were the same.

### Recent decrease in abundance of Cryptophytes in Lake Mendota

One particular example of marked change in abundances of eukaryotic communities in Lake Mendota is the decrease of abundance and diversity of cryptophytes. The overall diversity of cryptophytes in Lake Mendota is poorly characterized via microscopy, and may respond quickly to changes in the lake, based on studies of other similar temperate lakes [86]. In the NTL-LTER phytoplankton time series, only two genera of cryptophytes are documented [87], whereas our results suggest the presence of at least seven genera, and multiple putative cryptophyte MAGs. The lack of attention to cryptophyte diversity in previous studies could be because of methodological limitations in the microscopy used for the NTL-LTER measurements, as cryptophytes are smaller than many eukaryotic algae, and their size may also be variable *in situ* [88]. Cryptophyte abundance may be changing in response to other ecological shifts in Lake Mendota. Cryptophytes are known to be adapted to low-light conditions [89], so higher light intensity or smaller blooms of other organisms may reduce the available cryptophyte niche. Another potential factor is the presence of zebra mussels (*Dreissena polymorpha*), which are predicted to increase in abundance in response to increases in spiny water fleas [25]. The introduction of zebra mussels to Lake Oneida in the late 1980s was observed to correspond to a decrease in eukaryotic cryptophytes [86]. After 2015, a zebra mussel irruption was documented in Lake Mendota, with heretofore unknown impacts on the eukaryotic microbial community of the lake [90]. To our knowledge, no other study has approached recent changes in cryptophyte abundance, in particular with relation to spiny water flea and zebra mussel invasions. Taken together, our findings indicate that there has been a decline in cryptophyte diversity and abundance in Lake Mendota in recent (2010-2019) years, which may also have increased the available niche space for other eukaryotic organisms.

### Metagenome-assembled genomes reveal only the dominant fraction of Lake Mendota’s protistan diversity

Our results demonstrate that it is possible to link highly complete eukaryotic MAGs to an 18S rRNA gene sequence, and to use this to clarify the overall representation of assembled eukaryotic contigs in a mixed environmental metagenomic assembly.

The number of MAGs we recovered is on par with other recent studies from similar sample types [91], but we note that the complete MAGs we identified constitute a small fraction of the overall eukaryotic microbial diversity that we identified in the assembled contigs from the metagenomic samples via 18S rRNA gene discovery. Our observation that the vast majority of MAGs did not contain discovered 18S rRNA gene sequences from our discovery pipeline is in line with previous observations that these genes do not tend to co-bin with MAGs [92]. Occasionally, 18S rRNA gene sequences were not consistently extracted between samples. Hence although we did find a small subunit rRNA sequence in the dataset corresponding to a MAG, it was filtered out of the analysis. For example, while we could not directly attribute an OTU to *Bythotrephes longimanus* that met the threshold for use in the study (10 distinct occurrences via OTU clustering), we identified a single rRNA gene from the dataset that appears to be *Bythotrephes longimanus*, which was subsequently excluded due to not being present in multiple samples.

Successful bridging of more ubiquitous 18S rRNA gene amplicon studies and forthcoming extraction and characterization of eukaryotic MAGs from metagenomes depends on the ability to link and interpret these data types. We have shown in this study that MAGs and 18S rRNA gene sequences provide complementary insights into protistan diversity within a single metagenomic time series, and have the forthcoming potential to efficiently link multiple sources of -omic data. In particular, we envision that MAGs from samples enriched with a single organism will be a powerful tool for maximizing inference from time-series metagenomic surveys, because once MAGs are identified from the enriched samples, the putative genomes can be quantified across the entire time series. We did this for two enriched metagenomes collected for *Daphnia pulicaria* and *Bythotrephes longimanus*. While the MAGs extracted had relatively low completeness, this approach was far superior to the community-level metagenomes in allowing a MAG to be binned in the first place. Because count data suggest high abundances [93], we suspect metazoan sequences may be limited technically by assembly. Their low abundance may also be explained by whether full organisms were retained during the filtering process, in particular during intense blooms of gelatinous *Bythotrephes longimanus* that may have clogged the sample collection filters. If individuals were avoided, the only source of sequence data would be environmental DNA. However, abundance quantification both via the 18S rRNA gene extracted from the time series metagenomes and the extracted MAG from the enriched metagenome strongly suggest that *Bythotrephes longimanus* may have been present in the lake in low numbers before 2009 (Figure 5), which has been previously discussed in the literature [22, 94]. Past work in lake sediment cores suggests that the spiny water flea was present in low abundances in Lake Mendota since at least 1999 [23, 24] as a “sleeper population” until conditions were right for the organism to become dominant. Our analysis of this long-term time series provides additional evidence that suggest that the spiny water flea could be detected genetically in the lake prior to 2009 (5). Without collecting metagenomic time series in a similar way to the established method of long-term sediment coring, this powerful retrospective insight into the spiny water flea invasion and its effects would not have been possible. Enriched metagenomes and MAG binning for other organisms could yield more accurate estimation of their composition in the lake, and enable insights about the functional potential of taxa without sequenced genomes.

### Predicted eukaryote-prokaryote associations are common in Lake Mendota

Our analysis predicted a wide range of association between diverse bacteria and eukaryotes, in particular chlorophytes, diatoms, chytrid fungi, and ciliates.

We found correlations and putative associations between chlorophyte (green algal) OTUs and OTUs of several bacteria identified by other studies to be chlorophyte microbiome constituents, including families *Comamonadaceae*, *Rhodobacteraceae*, and *Chthoniobacteraceae* [95, 96], and *Flavobacteriaceae* and order *Sphingomonadales* (Alphaproteobacteria) [97]. Correlations were also identified between chlorophytes and sulfur bacteria (family *Chromatiaceae*) which are known to be abundant in lakes [98], and cyanobacteria of order *Oscillatoriales* (family *Microcoleaceae*), and two families of potentially pathogenic Gammaproteobacteria, *Moraxellaceae* and *Aeromonadaceae* [99, 100]. Chlorophytes are known to thrive under high nutrient concentrations early in the season, in particular high nitrate levels and a deep mixed layer [101, 102, 103]. This taxonomic group may increase with high levels of nutrient pollution, in particular extreme cases of nutrient pollution, alongside cyanobacteria [103, 102]. Chlorophytes are also known to have a microbiome of cooperative bacteria [95]. Hence, in our study we have simultaneously provided evidence that certain bacterial OTUs associate with chlorophytes in Lake Mendota, as well as identified OTUs of bacteria and may similarly increase in concert with nutrient pollution in conjunction with chlorophytes [103].

We found a diatom (phylum *Ochrophyta*, class *Bacillariophyta*) of order *Aulacoseirales* that was significantly correlated several other bacterial and eukaryotic OTUs: one ciliate, a bacterium of order *Spirochaetales* which is known to be present among some diatom microbiota [96], cyanobacteria of orders *Chroococcales* and *Synechococcales*, one OTU of order *Verrucomicrobiales*, and an Alphaproteobacterium of order *Rickettsiales*. Several of the observed bacterial associations, in particular cyanobacteria of orders *Chroococcales* and *Synechococcales*, may be related to their shade-tolerance during dense blooms of diatoms and other eukaryotes [104, 105]. A second diatom only had a strong, significant correlation to *Synechococcales* (*Chamaesiphonaceae*), but not only was this a distinct OTU as indicated by our clustering approach, it was one of the strongest correlations we recorded in the dataset. This may be related to the observation that *Chamaesiphonaceae* is a major component of some biofilms [81], and diatom frustules have been observed to be embedded in biofilms present in other lakes [106, 107]. By contrast to diatoms, a dinoflagellate OTU of order *Gymnodiniales* predicted to be associated with *Desulfuromonadales*, *Geodermatophilales*, and *Acidimicrobiales*, organisms that tend to respond strongly to availability of excess organic matter [108], or may have uncommon defenses against oxidative stress [109]. Two other bacterial orders found to be linked to a dinoflagellate OTU of order *Peridiniales*, *Saprospirales* and *Sphingomonadales*, were found to be associated with phosphate depletion and high phosphate scavenging in a previous laboratory study [110]. Notably, the *Peridiniales* OTU significantly correlated with an cercozoan and a ciliate OTU, and these dinoflagellates are known to be mixotrophs that may feed on ciliates and other large protists [111]. Diatoms are associated with high nutrients and turbulent water column conditions, and tend to have highest abundance earlier in the spring than the other protistan taxa that we considered [101, 102]. In ocean ecosystems replacement of diatoms with dinoflagellates has been observed as a potential response to rapid excess nutrient loading [112]. Hence, these associations may reflect differing trophic preferences for dinoflagellates as they inhabit novel niches under nutrient stress and changing community composition.

The analysis also suggested an association of Chytridiomycota and planctomycetes of the order Isosphaerales. Both *Chytridiomycota* and planctomycetes are known to be degraders of organic matter in lake ecosystems [113, 114, 115]. Hence, these two taxa may be involved in cleanup of organic matter in the lake. The greatest abundances of *Chytridiomycota* and *Isosphaerales* were observed in April, which might be associated with the influx of organic matter observed with spring runoff during the ice-off period in lakes [116]. Due to uneven sampling during the winter period at the beginning of the time series, it remains unknown whether earlier ice-off due to increasing temperatures is changing the prevalence of *Chytridiomycota* and *Planctomycetes*. Both taxa likely play a role in the degradation of organic matter, but there is some evidence of their increased abundance (Supplementary Figure 2). Interestingly, some other *Planctomycetes* showed correlations only with individual eukaryotic OTUs or distinct taxonomic groups, which may indicate that they depend on the presence of eukaryotic taxa as specialized hosts [117]. Planctomycetes are also known to use the nutrients generated from algal or cyanobacterial blooms as a food source [117], which might explain these observed correlations. Of particular note is the strong correlation between a dinoflagellate and a planctomycete, which may support the hypothesis that specific bloom types offer substrates for these bacteria [118, 117].

Our study revealed a broad diversity of ciliates that formed a high number of clusters with both bacteria and other eukaryotes. These results are particularly informative because many species of ciliates, apicomplexans, and cryptophytes can be quite challenging to delineate based on morphological features, despite their abundance in the lake system [119, 120]. Among the putative interactions, there was one coherent module that consisted of two ciliates (families *Vorticellidae* and *Acinetidae*) and a *Bacteroidetes* OTU of family *Sphingobacteriaceae*. This strong association may be related to an endosymbiotic relationship between the bacterium and one or both of these ciliates, as a *Sphingobacteria* has previously been shown to have an endosymbiotic relationship with *Ichthyophthirius multifiliis* ([121]), an organism in the same taxonomic class as one of these ciliates (*Oligohymenophorea*). Some bacteria are associated with ciliates because of being resistant to digestion by the ciliate [122]. We identified at least one of these bacteria, family *Phycisphaeraceae*, that clustered with a ciliate OTU, highlighting yet another route by which eukaryotes may be associated with bacteria in this system.

### Long term time series facilitated the detection of invasive species responses and putative associations, which provides a foundation for future testable hypotheses

The prediction of protistan relationships in long-term time series metagenomic data from Lake Mendota supports and extends our understanding of the roles played by the major taxonomic groups in lake ecosystems and their effects on food web structure [123]. We identified putative interactions between protists and bacteria on the basis of their consistently correlated abundances, which opens the door for exploration of novel interactions between these taxa, including lifestyles like endo- or ectosymbioses [124] or farming (whereby organisms like amoebas may tote and proliferate their bacterial food) [125]. Our results provide evidence for the presence and abundance of such interactions, and enable new perspectives on their ecology. Further, our metagenomic identification of a possible “sleeper population” of spiny water flea aligns with observations from previous research [22, 24], provides further genetic contextualization for this important invasion event, and highlights the importance of long-term ecosystem surveillance using metagenomics.

**Table 1:**
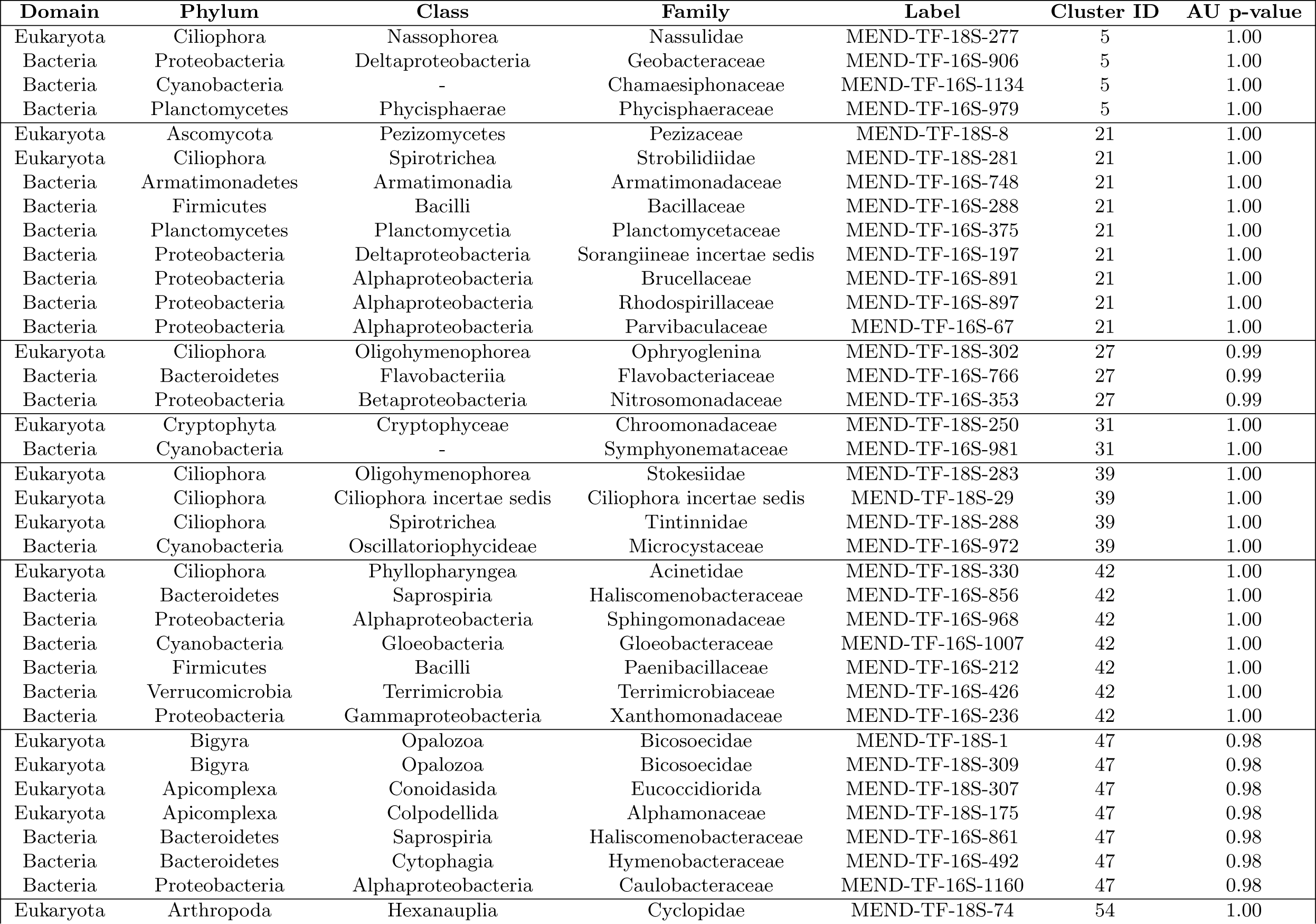
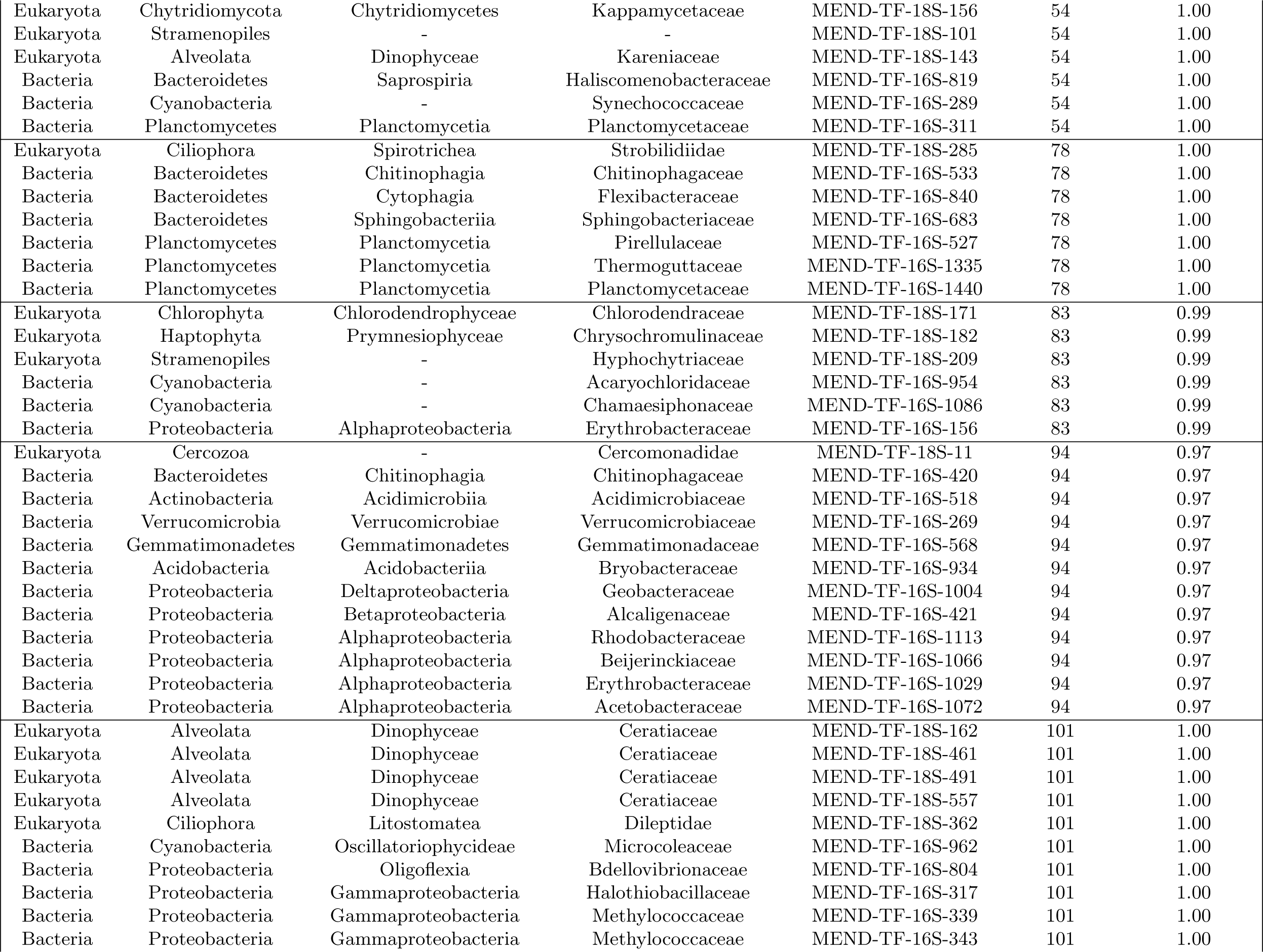
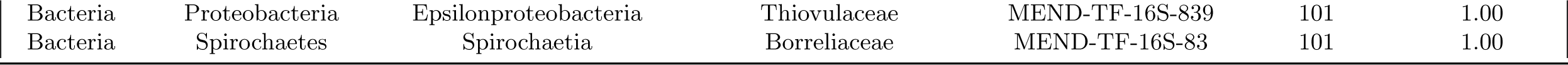
List of extracted SSU rRNA genes from the Lake Mendota metagenomes and the hierarchical cluster that they were contained in.

## Acknowledgements

The authors gratefully acknowledge members of the McMahon Lab at the University of Wisconsin-Madison for graciously sharing existing data and expertise, and AIK’s thesis advisors, Dr. Michael Follows and Dr. Harriet Alexander, for their support and feedback during the completion of the work. The authors also sincerely thank the curators of the North Temperate Lakes time series for access to phytoplankton and zooplankton count data from Lake Mendota.

## Abbreviations

rRNA: ribosomal RNA gene
MAG: metagenome-assembled genome

## Availability of data and materials

The processed dataset and accompanying code supporting the conclusions of this article are available in the akrinos/2022-Krinos-Mendota18s repository, https://github.com/akrinos/2022-krinos-mendota18s.

The original TYMEFLIES metagenomes from which 18S sequences were originally extracted are available from the Joint Genome Institute (JGI; https://genome.jgi.doe.gov/portal/Exttemetagenomes/Exttemetagenomes.info.html); the list of project and sample numbers which were used, including the assembly products produced by the JGI’s automated analysis pipeline is available in Supplementary Table 1.

The extracted 16S and 18S rRNA genes as well as the eukaryotics metagenome-assembled genomes (MAGs) found to be highly complete are available via the Open Science Framework (OSF) at this link: https://osf.io/9epa8/?viewonly=152af26e11894ac0bcdfe542e02c6ab1.

## Authors’ contributions

AIK, FS, TW, and RMB designed the co-occurrence study. AIK and FS refined methods for rRNA gene extraction. AIK processed and analyzed data and generated figures and statistical analyses. KDM oversaw original data collection, provided feedback and ecological insight, and helped refine text. AIK wrote the paper with editing and input from all authors.

