## Supplemental Text and Figures for "Time-series metagenomics reveals changing protistan ecology of a temperate dimictic lake"

### Supplementary Information

#### Supplemental Introductory Material

##### General Description of Lake Mendota

Lake Mendota, Dane County, Madison, WI, USA is a large (3961 ha in area, 25 m at maximum depth) dimictic lake which provides ecosystem services to its neighboring city and state [11]. Because of the ecological relevance of Lake Mendota, consistent physicochemical measurements are made at the site as part of the North Temperate Lakes Long-Term Ecological Research Program. Microbial eukaryotes in Lake Mendota tend to have highest abundance in the spring and fall [126, 127, 128]. These cells tend to be larger in size than the smaller bacterial cells that dominate during summer blooms, and eukaryotic blooms in the springtime may be diatom or dinoflagellate-majority [11]. Either zooplankton like genus *Daphnia* consume the eukaryotic phytoplankton or nutrient limitation terminates the bloom, resulting in a "clear-water phase" that makes way for summer cyanobacteria [11].

Recent ecological intervention and invasion in Lake Mendota has complicated the food web dynamics in the system that microbial eukaryotes inhabit. In particular, stocking of Lake Mendota with fish species like walleye and pike reduced populations of zooplanktivorous species like cisco [129, 22]. This led to an increase in populations of larger *Daphnia pulicaria* zooplankton in relation to smaller *Daphnia mendotae* populations, which helped to keep water quality in check, as the larger zooplankton were more capable of consuming larger amounts of the algae that impair water quality [129]. More recently, an invasive predator, the spiny water flea *Bythotrephes longimanus* (hereafter, *Bythotrephes*) has been able to establish itself in Lake Mendota, changing the zooplankton population. Namely, larger cladocerans of species *Daphnia pulicaria* have declined in relation to smaller predation-resistant *Daphnia mendotae*, reducing water quality and increasing the magnitude of blooms of eukaryotic phytoplankton [84]. Despite the significance of the establishment of the *Bythotrephes* invasion, the change went initially undetected, and continues to have unseen consequences on ecosystem functioning [23] and hence predation pressure on microbial eukaryotes [85].

Cyanobacterial blooms are known to be tightly coupled to the abundance of Lake Mendota's protists. Some cryptophytes may ingest the toxic cyanobacterium *Microcystis aeruginosa*, reducing toxin levels in lakes during cyanobacterial blooms [130]. Eukaryotes and bacteria may coexist as colonizers of lake macroalgae [10]. A year of measurements of Lake Texoma (Texas/Oklahoma, U.S.A.) showed ecophysiological relationships between a toxic alga and bacteria and other microbial eukaryotes [131], and high-frequency flow cytometric measurements of bacteria and eukaryotes clarified the trade-off between light and temperature conditions which may enable co-existence between microbial eukaryotes and fast-growing bacterial taxa [132]. Paleoclimactic analysis of past patterns of microbial eukaryotes can be reconstructed that reveal changes in the abundance of certain eukaryotic microbes and widespread homogenization in freshwater ecosystems [20], and sedimentary records have also shown the sensitivity of microbial eukaryotes to environmental conditions, in particular of alveolates to temperature [133]. Time series analysis is a much more mature field when it comes to Lake Mendota's bacteria, in particular its large seasonal cyanobacterial blooms, which are the consequence of persistent nutrient pollution [11] and are generally increasing due to climate change [134, 135, 136, 137]. Ribosomal intergenic spacer analysis has been applied to bacterial populations in Lake Mendota to prove the strong seasonality of bloom networks [138]. Recent metatranscriptomic analysis has proven the diel character of gene expression of photosynthesis and sugar transport in phytoplankton communities in Lake Mendota over a two-day time series, but taxonomic analysis identified only bacterial representatives [139]. Time-series metagenomic analyses and single-cell genomics have recently exposed the presence of genetically distinct "tribes" within bacterial lineages, in particular the acl clade within the Actinobacteria [140], and clarified the taxonomy and ecophysiology of *Verrucomicrobia* [141].

The same time-series metagenomes from Lake Mendota have also been used to identify new giant virus lineages that showed highest diversity in 2012 [142], after cyanobacterial populations first peaked in 2008 due to nutrient input from particularly strong spring floods [143], and then spiny water flea invasion reduced the abundance of predators of protists [22]. The authors of this study searched the lake metagenomes for 18S rRNA gene sequences, but only over a five-year time period [142]. They explore eukaryotes within the groups *Dinophyceae*, *Rhizaria*, *Alveolata*, and *Cryptophyceae* as potential giant virus hosts, but do not attempt to define the taxonomy of these sequences beyond the order level, nor explain further implications of their changes in abundance apart from the correlation of these changes to changes in giant virus abundance.

### Supplementary Methods

#### Use of the SILVA and PR2 databases

We also identified the most likely rRNA candidate from among the SILVA [144, 145, 146] and PR2 [147, 148] databases using the `DIAMOND` [149] alignment tool, prioritizing the more comprehensive PR2 database for eukaryotic hits, and using the eight-part taxonomic labeling system for eukaryotes as organized in PR2 [147]. Hits with an e-value below 1e-5 were retained, and further the best match on the basis of percentage identity was selected, with a maximum of two sequences selected per contig, one per rRNA model.

#### Comparison of rRNA gene identification to observations

Since 1995, consistent measurements of the phytoplankton community in Lake Mendota have been collected at the Deep Hole site as part of the North Temperate Lakes Long-Term Ecological Research (LTER) program [87, 150]. We used these counts for several major taxonomic groups of interest to validate the general trends found in the 16S and 18S gene-derived normalized abundance estimates. Similar to the procedure used for the rRNA gene abundances, we calculated a Z-score for the count data expressed in cells per milliliter. Although Z-scores were calculated across entire subset periods of interest for co-occurrence relationships and community identification in subsequent chapters, Z-scores were calculated within years for the purposes of Pearson correlation for groups that showed significant changes in average abundance over the course of the 20-year time series, so as to emphasize the identification of blooms and to avoid bias due to low abundance years or those missing multiple sampling points. We used linear imputation to correct for points missing in either dataset according to the nearest calendar week, such that a Pearson correlation could be calculated between the time series for the period from 2000 to 2018 during which count data are available. Linear interpolation was performed using `SciPy` [151], and Pearson correlation was calculated using the `pandas` library [152]. All analyses were conducted in Python version 3.7 [153] or R version 4.1 [43].

### Supplementary Results

#### rRNA discovered from Lake Mendota metagenomes approximates trends from the North Temperate Lakes Time Series

Pearson $R^2$ correlations between time series of Z-scores from rRNA abundance estimates derived from metagenomes and phytoplankton count data from the NTL LTER site at Mendota were high for several major and abundant groups of organisms. Z-scores for the second half of the time series, during which period metagenomes were collected more frequently, Pearson $R^2$ values tended to be higher. Pearson $R^2$ values expressed the match in the trend between the North Temperate Lakes (NTL) time series and the extracted rRNA gene sequences, while linear regression values calculated between the two time series tended to be lower. Nevertheless, the more specific the taxonomic annotation and the more precise and exclusive the OTU, the better the correlation between the time series tended to be. In particular, family *Fragilariaceae* had a Pearson correlation of 0.827 ($p < 2.2e^{-16}$) between the extracted rRNA and the NTL time series (linear regression $R^2 = 0.683$, $p < 2.2e^{-16}$), order *Prymnesiales* had a Pearson correlation of 0.585, and order *Aulacoseirales* had a Pearson correlation of 0.515 ($p < 4.6e^{-16}$). This was despite the fact that class *Coscinodiscophyceae*, one taxonomic level above order *Aulacoseirales*, had a Pearson correlation of only 0.220 ($p = 0.0011$).

Some groups of organisms were notably difficult to correlate to the metagenomic abundance estimates. In particular, cryptophytes were difficult to compare between the North Temperate Lakes time series and the Mendota metagenomes. Only two genera of cryptophytes were present in the North Temperate Lakes time series: *Cryptomonas* and *Rhodomonas*, and a single species *Rhodomonas minuta*, showed significant increases in the later half of the time series (Supplementary Figure 4).

#### Changes in bacterial community composition

Some bacterial taxa showed similar increases in abundance after 2010, although unlike eukaryotes, some bacterial taxa also showed significant decreases in abundance after 2010 as compared to the first half of the time series. While 277 identified bacterial OTUs extracted from at least 10 metagenomes did not show significant change from the first half of the time series to the second, 124 showed significant decreases, and 207 showed a detectable increase according to the same p-value adjustment procedure as given above.

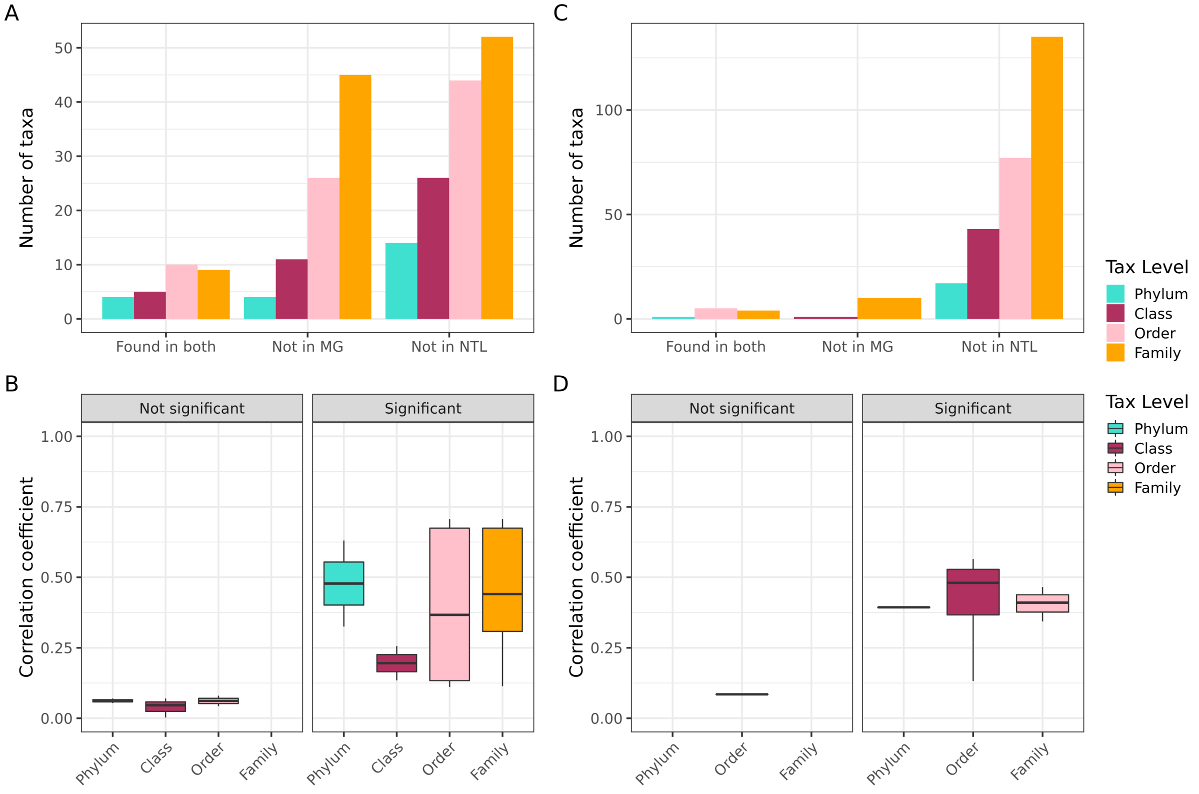


Figure 1: Graphical summary of the extent to which comparisons could be made between the North Temperate Lakes dataset and the extracted 16S and 18S rRNA genes from the time series metagenomes. A: For eukaryotes, the number of taxa that were (leftmost group of bars) found in both the NTL and metagenomic datasets, (middle group of bars) found in the North Temperate Lakes dataset but not the metagenomic dataset, and (rightmost group of bars) found in the metagenomic dataset but not the North Temperate Lakes dataset. B: For eukaryotes, a boxplot of correlation coefficients for (left) correlations that were not statistically significant and (right) correlations that were statistically significant for each taxonomic level. C: The same as panel A, but for bacteria. D: The same as panel B, but for bacteria. The preponderance of bacteria were not part of the phytoplankton dataset, and similarly many metazoans within the eukaryote dataset extracted from the metagenomes were not found in the North Temperate Lakes data collection. However, many eukaryotes were also recorded in the North Temperate Lakes dataset but did not correspond to an 18S rRNA gene OTU, likely due to a combination of limitations in morphological identification and errors in identifying the correct annotation for the extracted 18S rRNA gene.

#### Validation with North Temperate Lakes Time Series

Several taxa of interest performed impressively well with respect to Pearson correlation and linear regression between the North Temperate Lakes time series and the extracted values from the metagenomes. Although the Pearson correlations were not high for all groups, this can be attributed to multiple limiting factors preventing fair comparison of the two datasets. Firstly, the use of the Z-score is an improvement over absolute values, but an imperfect metric. Secondly, several of the organism we were comparing are notoriously difficult to identify or distinguish microscopically, leading to uncertainties in the microscopic counts relative to the taxonomic annotation derived from the metagenomic rRNA. Additionally, count data was not available for the same timepoints as the metagenomes

| Taxon | Taxon Level | | Pearson Correlation Coefficient | Pearson $p$-value | Domain |
|---|---|---|---|---|---|
| Cyanobacteria | Phylum | Found in both | 0.39 | 1.11e-19 | Bacteria |
| Synechococcales | Order | Found in both | 0.57 | 6.70e-43 | Bacteria |
| Nostocales | Order | Found in both | 0.44 | 2.05e-24 | Bacteria |
| Pseudanabaenales | Order | Found in both | 0.52 | 5.26e-10 | Bacteria |
| Oscillatoriales | Order | Found in both | 0.13 | 7.30e-03 | Bacteria |
| Chroococcales | Order | Found in both | 0.08 | 5.98e-02 | Bacteria |
| Microcystaceae | Family | Found in both | 0.41 | 5.84e-19 | Bacteria |
| Microcoleaceae | Family | Found in both | 0.47 | 1.53e-14 | Bacteria |
| Nostocaceae | Family | Found in both | 0.34 | 1.13e-11 | Bacteria |
| Synechococcaceae | Family | Found in both | -0.12 | 8.82e-03 | Bacteria |
| Haptophyta | Phylum | Found in both | 0.63 | 8.18e-49 | Eukaryota |
| Ochrophyta | Phylum | Found in both | 0.33 | 5.67e-13 | Eukaryota |
| Cryptophyta | Phylum | Found in both | 0.07 | 1.21e-01 | Eukaryota |
| Chlorophyta | Phylum | Found in both | 0.05 | 2.34e-01 | Eukaryota |
| Coscinodiscophyceae | Class | Found in both | 0.26 | 5.28e-07 | Eukaryota |
| Dinophyceae | Class | Found in both | 0.13 | 9.24e-03 | Eukaryota |
| Cryptophyceae | Class | Found in both | 0.07 | 1.21e-01 | Eukaryota |
| Chrysophyceae | Class | Found in both | 0.05 | 3.64e-01 | Eukaryota |
| Chlorophyceae | Class | Found in both | 0.00 | 9.54e-01 | Eukaryota |
| Fragilariales | Order | Found in both | 0.69 | 1.17e-52 | Eukaryota |
| Prymnesiales | Order | Found in both | 0.66 | 3.78e-27 | Eukaryota |
| Gonyaulacales | Order | Found in both | 0.71 | 5.37e-26 | Eukaryota |
| Aulacoseirales | Order | Found in both | 0.37 | 2.89e-13 | Eukaryota |
| Pyrenomonadales | Order | Found in both | 0.14 | 2.59e-03 | Eukaryota |
| Peridiniales | Order | Found in both | 0.13 | 1.92e-02 | Eukaryota |
| Sphaeropleales | Order | Found in both | -0.10 | 2.92e-02 | Eukaryota |
| Gymnodiniales | Order | Found in both | 0.11 | 3.32e-02 | Eukaryota |
| Chlamydomonadales | Order | Found in both | 0.08 | 8.13e-02 | Eukaryota |
| Chromulinales | Order | Found in both | 0.04 | 4.04e-01 | Eukaryota |
| Fragilariaceae | Family | Found in both | 0.69 | 1.17e-52 | Eukaryota |
| Chrysochromulinaceae | Family | Found in both | 0.66 | 3.78e-27 | Eukaryota |
| Ceratiaceae | Family | Found in both | 0.71 | 5.37e-26 | Eukaryota |
| Chlamydomonadaceae | Family | Found in both | 0.44 | 1.62e-22 | Eukaryota |
| Stephanodiscaceae | Family | Found in both | 0.31 | 5.48e-11 | Eukaryota |
| Volvocaceae | Family | Found in both | 0.30 | 1.54e-07 | Eukaryota |
| Cryptomonadaceae | Family | Found in both | 0.11 | 1.37e-02 | Eukaryota |
| Pyrenomonadaceae | Family | Found in both | -0.04 | 4.22e-01 | Eukaryota |
| Chromulinaceae | Family | Found in both | -0.03 | 6.21e-01 | Eukaryota |

Table 2: Pearson correlations for a selection of taxa from the extracted rRNA gene sequences from the Lake Mendota TYMEFLIES dataset as compared to the count data from the North Temperate Lakes timeseries. Taxa that were assessed at a more precise taxonomic level tended to have higher correlations, and several major taxa did not have annotations present in the North Temperate Lakes count data. In particular, alveolates and cryptophytes were underrepresented in the dataset despite evidence that they are present in Lake Mendota. Completely excluded from this analysis are non-algal bacteria that were not counted as part of the long-term monitoring effort.

in many cases (usually they were offset by some length of time). Because increases in abundance can be quite ephemeral, this could also lead to mismatch. The precise location, depth integration, and time of day of the two collections might also lead to differences in the abundance. The ability of the metagenomic rRNA to capture major maxima in abundance (Figure 2) is encouraging. Further, the improvement of the correlation when more precise taxonomic annotations were used in several cases is also encouraging, as it means that differences in labeling may be responsible for discrepancies. Further, we excluded rRNAs from the analysis and correlations if they were initially extracted from fewer than 10 samples per OTU clustering. This may be responsible for cases in which the estimated metagenomic abundance falls short of the reported value in the North Temperate Lakes time series, in particular for higher taxonomic levels which may encompass less abundant taxa. The fact that organisms of interest to the ecology of Lake Mendota (in particular, cyanobacteria and some diatoms) had high correlation performance is a good metric that the metagenomic rRNAs accurately represent lake dynamics, as these organisms have higher abundance and would be of both higher abundance and higher importance for count reporting.

### Supplementary Figures

### Supplementary Tables

#### Additional file 1 — Sequencing metadata

Information on dates of sampling, raw reads sequenced, and contigs assembled which were used in this study.

#### Additional file 2 — Hierarchical clustering results

Correlation coefficients and $p$-values for correlations between protists and bacteria.

#### Additional file 3 — Network analysis results

Network relationships predicted between taxa, their $p$-values and correlation coefficients, and the predicted taxonomy of the SSU rRNA gene sequences.

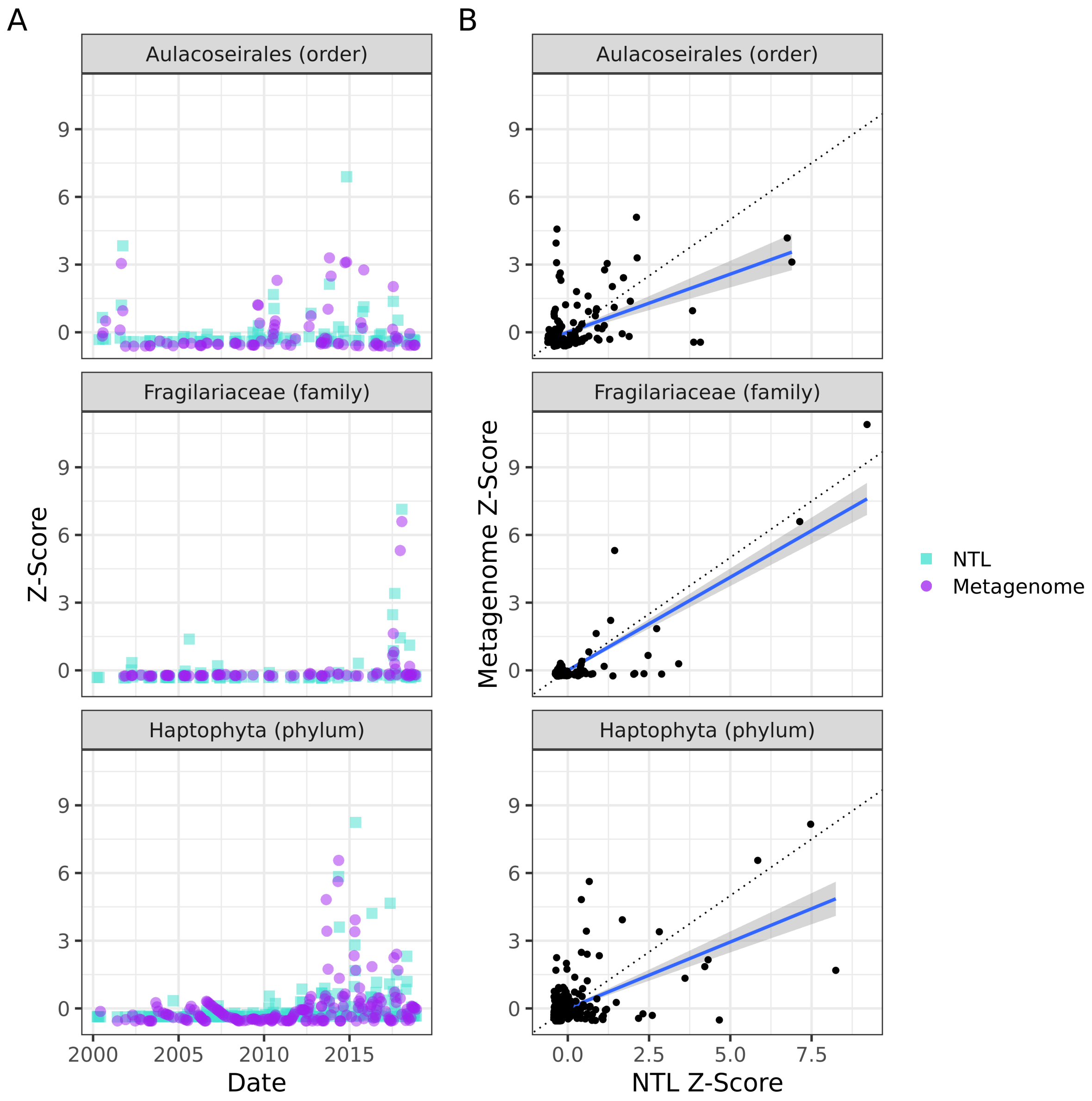


Figure 2: Comparison of three groups at the phylum, order, and family level. A: Z-scores over the full time series for each group as compared between the North Temperate Lakes count data (blue squares) and the extracted 16S/18S rRNA gene sequences from the TYMEFLIES metagenomes (purple circles). B: Comparison of the Z-score in the NTL timeseries (x-axis) to in the Mendota metagenomes (y-axis). The blue regression line shows the linear regression between the two sets of points with standard error, while the dotted line shows a one-to-one relationship.

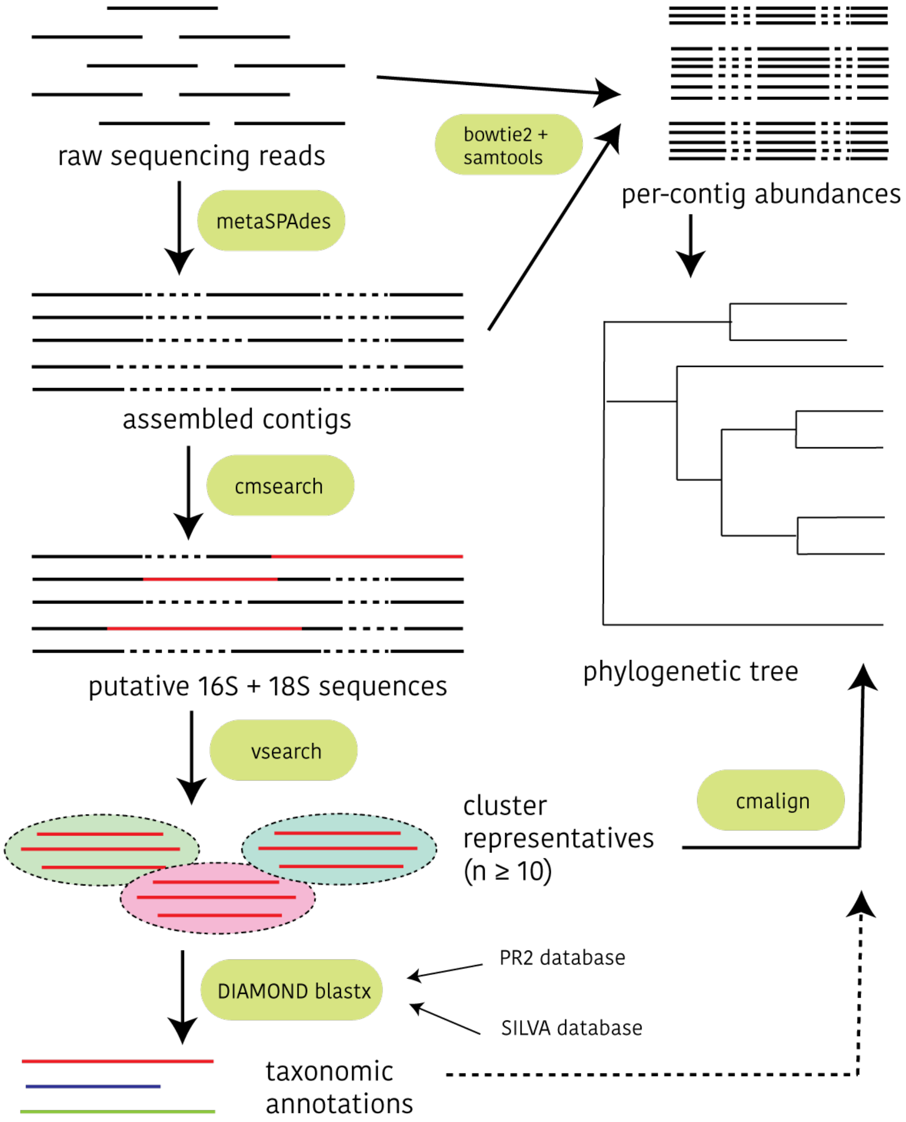


Figure 1: Workflow for extracting communities from Mendota metagenomes. In brief, raw sequencing reads were assembled into contigs, from which 16S and 18S rRNA gene sequences were extracted using a covariance model and the software `cmsearch` [31]. Subsequently, abundance information was calculated and a phylogenetic tree constructed in order to evaluate the placement of the extracted sequences and to validate taxonomic annotations created from a `DIAMOND` search against the PR2 and Silva databases [145].

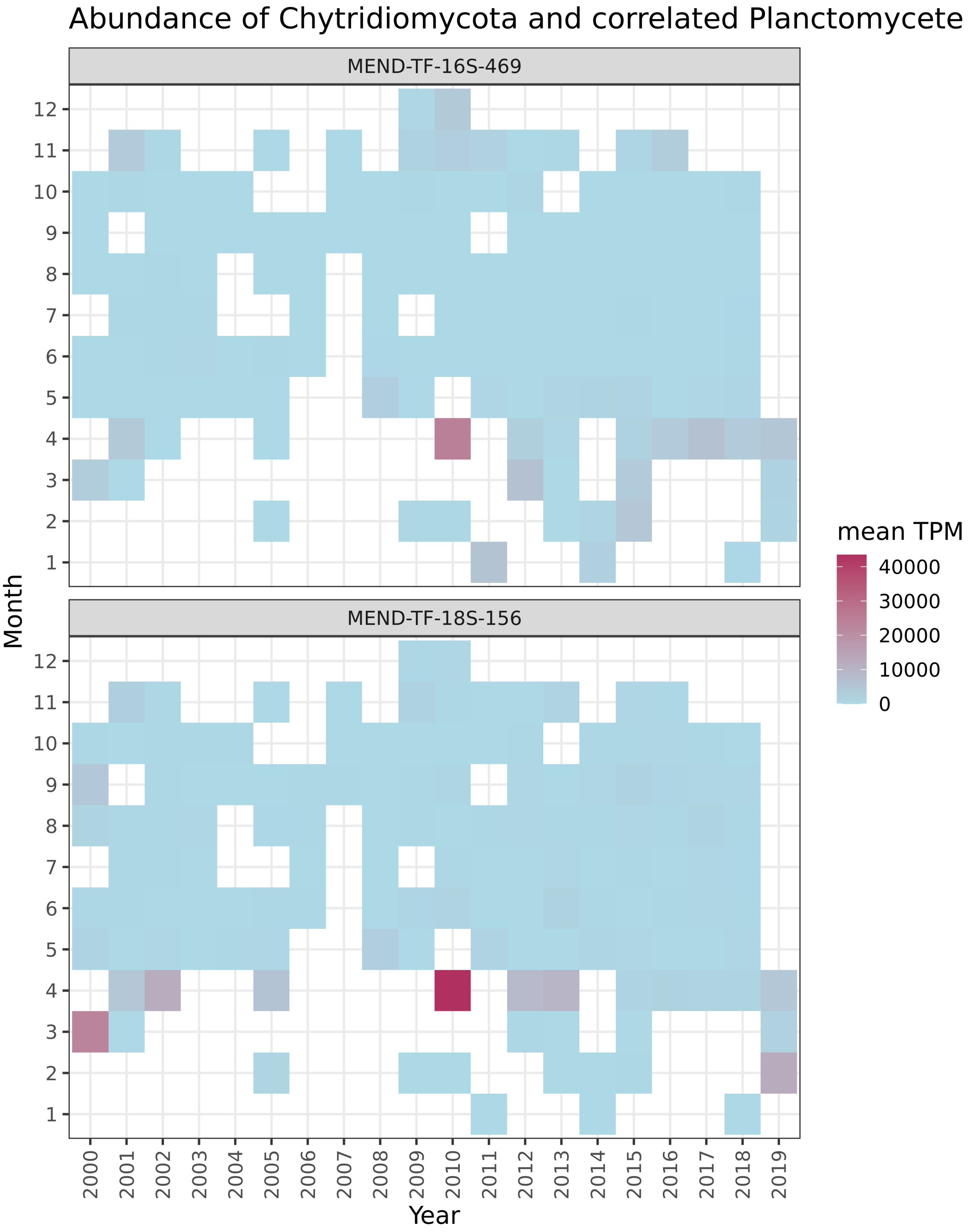


Figure 2: Abundance of *Chytridiomycota* OTU and correlated *Planctomycete* OTU aggregated over mean abundance by month and year in the time series metagenomes. Even though data were more sparse in the earlier part of the time series with respect to winter sampling, there is some evidence for increased early-year abundance of these organisms, or potentially for increasing abundance due to more rapidly melting ice.

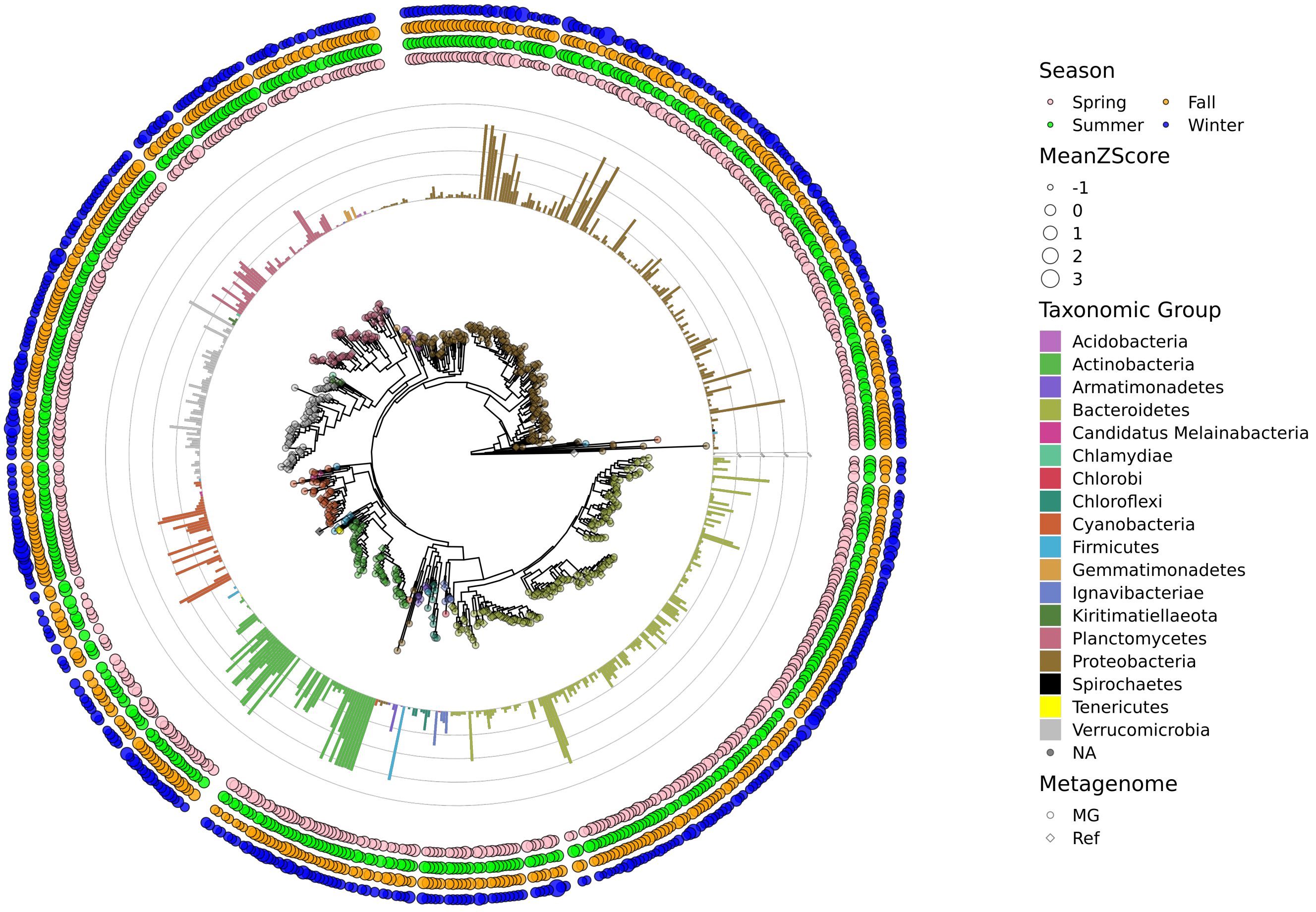


Figure 3: Phylogenetic tree for bacterial 16S sequences extracted from Lake Mendota metagenomes as part of the TYMEFLIES project. Bar plots show the number of extracted 16S sequences across all timepoints in each cluster, while the other circles are colored by season and sized proportionally to the average Z-score of each clustered OTU per season.

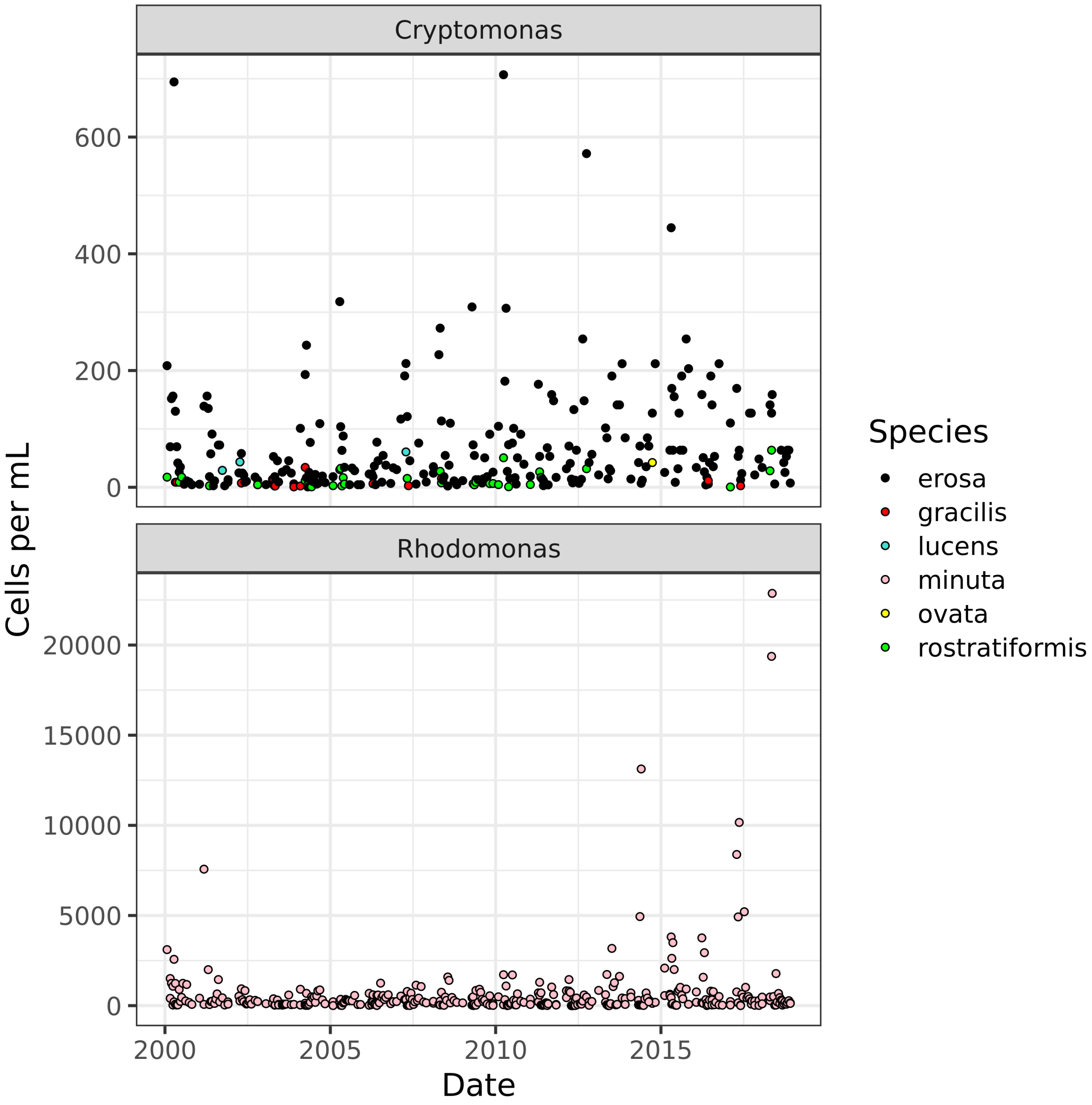


Figure 4: Cryptophytes in the North Temperate Lakes time series at the epilimnion site of Lake Mendota as measured in cells per mL of water. Genus *Rhodomonas* had a single species, *Rhodomonas minuta*, that increased significantly in abundance during the time series, whereas genus *Cryptomonas* had generally lower abundances.

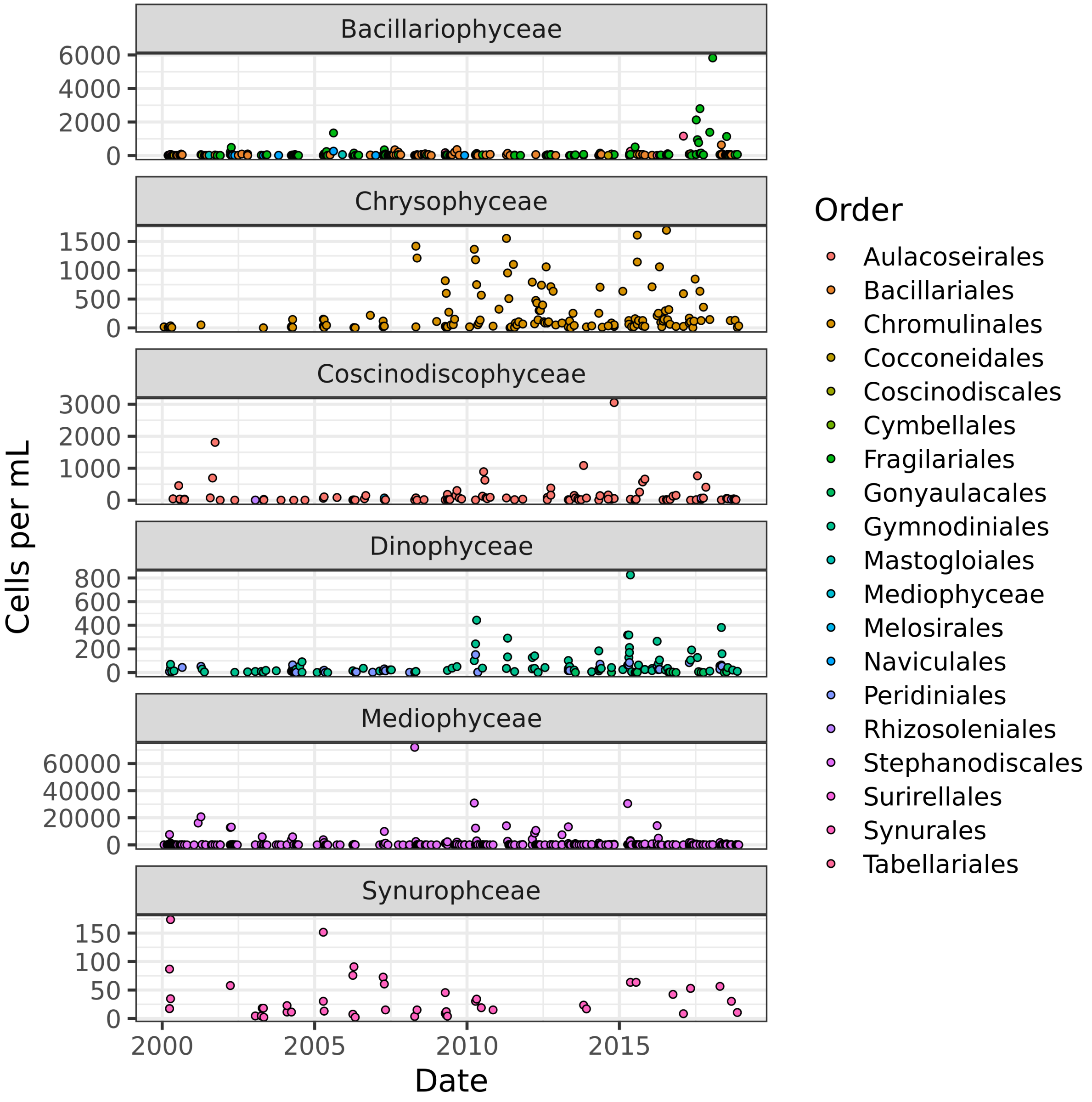


Figure 5: Dinoflagellates and ochrophytes in the North Temperate Lakes time series at the epilimnion site of Lake Mendota as measured in cells per mL of water. Only dinoflagellates were measured in the North Temperate Lakes time series; ciliates and apicomplexans were not reported.

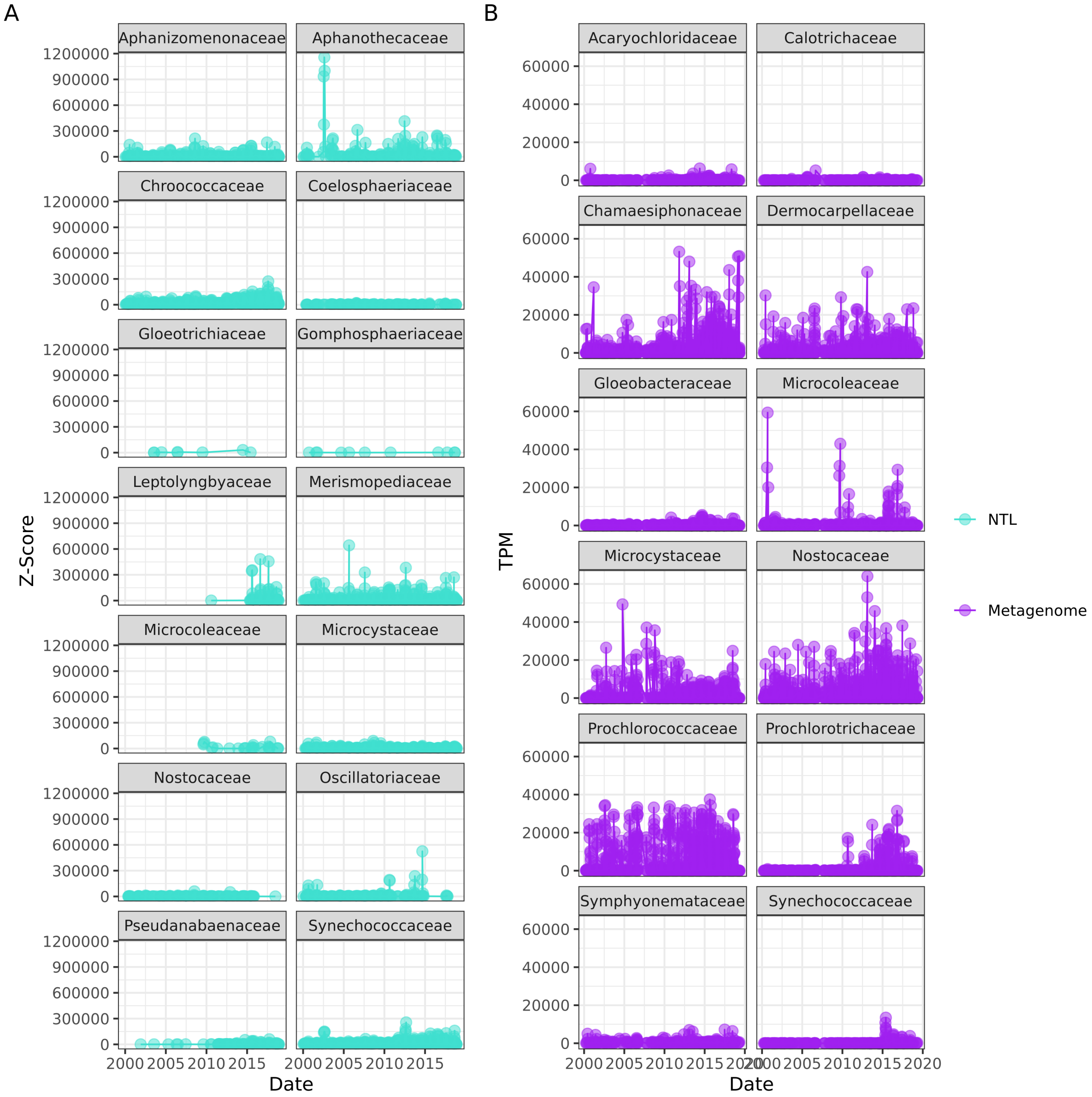


Figure 6: Raw abundance values for A: North Temperate Lakes count data in cells per mL and B: OTUs from Lake Mendota metagenomes (TYMEFLIES) with 10 or more occurrences in the dataset in estimated abundances for taxonomic families of phylum Cyanobacteria from the quantification procedure described in the methods section. These values are not normalized in order to show which groups were abundant in raw form. While some of the OTUs from the NTL dataset were extracted from the Mendota metagenomes, they may be missing from panel B due to low frequency of extraction. Some taxa, if recovered and annotated correctly in the Mendota metagenomes, would not appear in the NTL time series due to being too small to count.

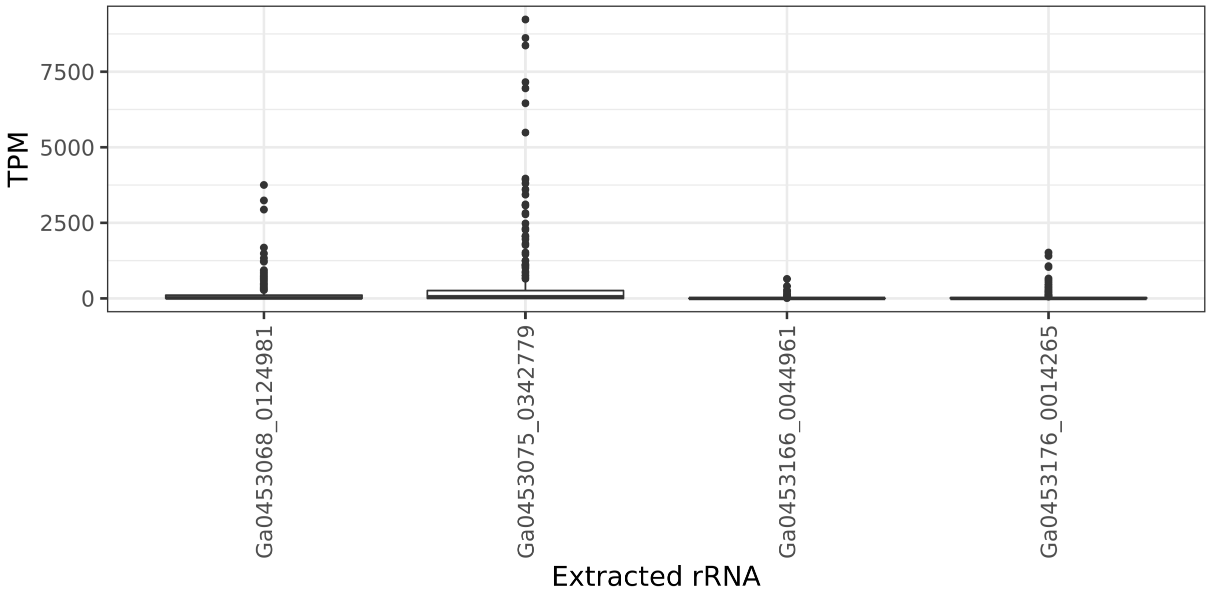


Figure 7: OTUs extracted from Mendota metagenomes of taxonomic Family *Aphanothecaceae* and their abundance across all data points. Despite a recorded bloom of this taxonomic family in the North Temperate Lakes count data, the extracted OTUs were not consistently recovered or found to be highly abundant, so were excluded from the lake profile created from the metagenomes.

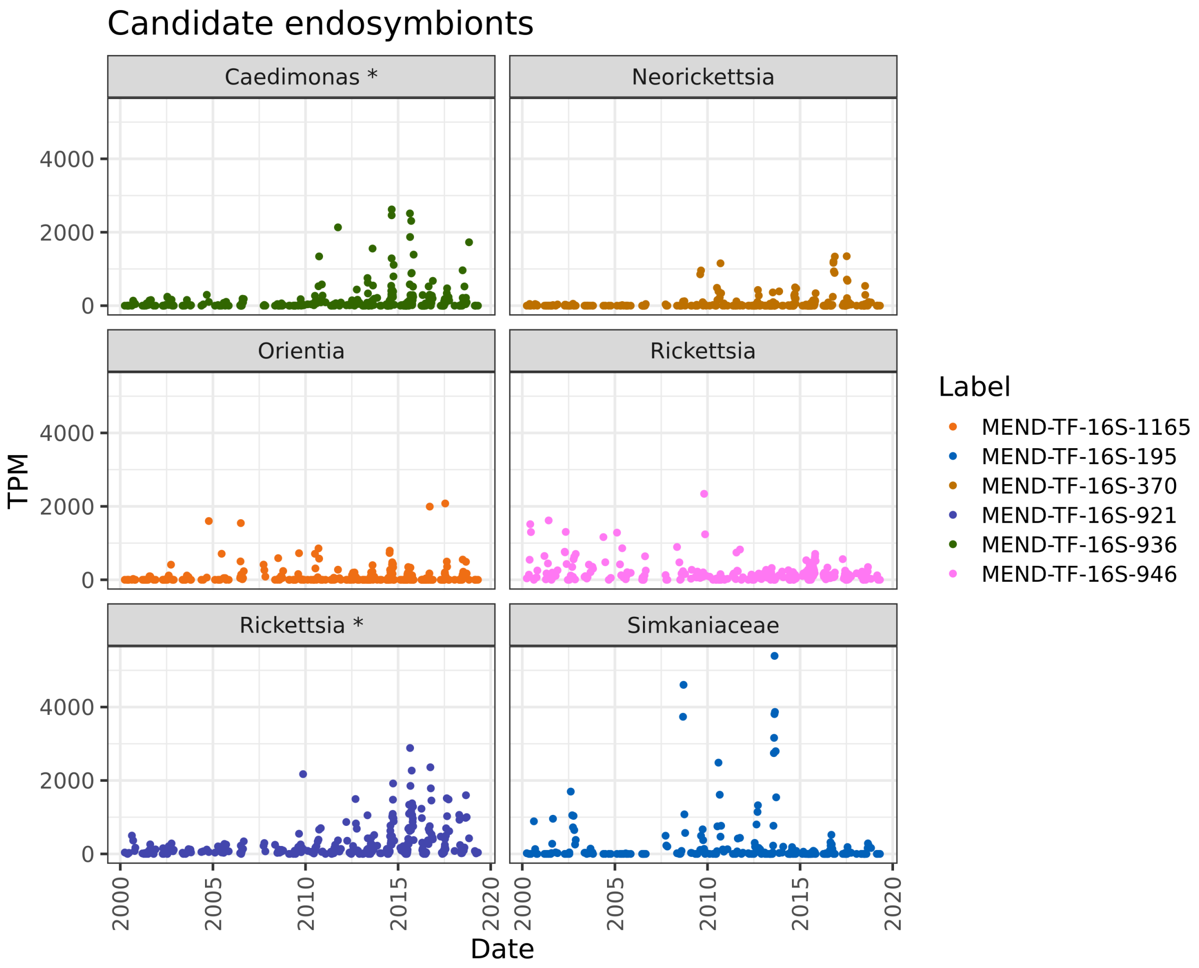


Figure 8: Time series of normalized abundances for genera of bacteria of phylum *Chlamydiae*, order *Rickettsiales* and order *Holosporales* recovered from the Mendota metagenomes that meet the threshold of 10 distinct sample identifications. Asterisks on facetted genus labels indicate that the adjusted $p$-value of the test of whether the abundance of the OTU increased significantly after 2010 was less than 0.05.

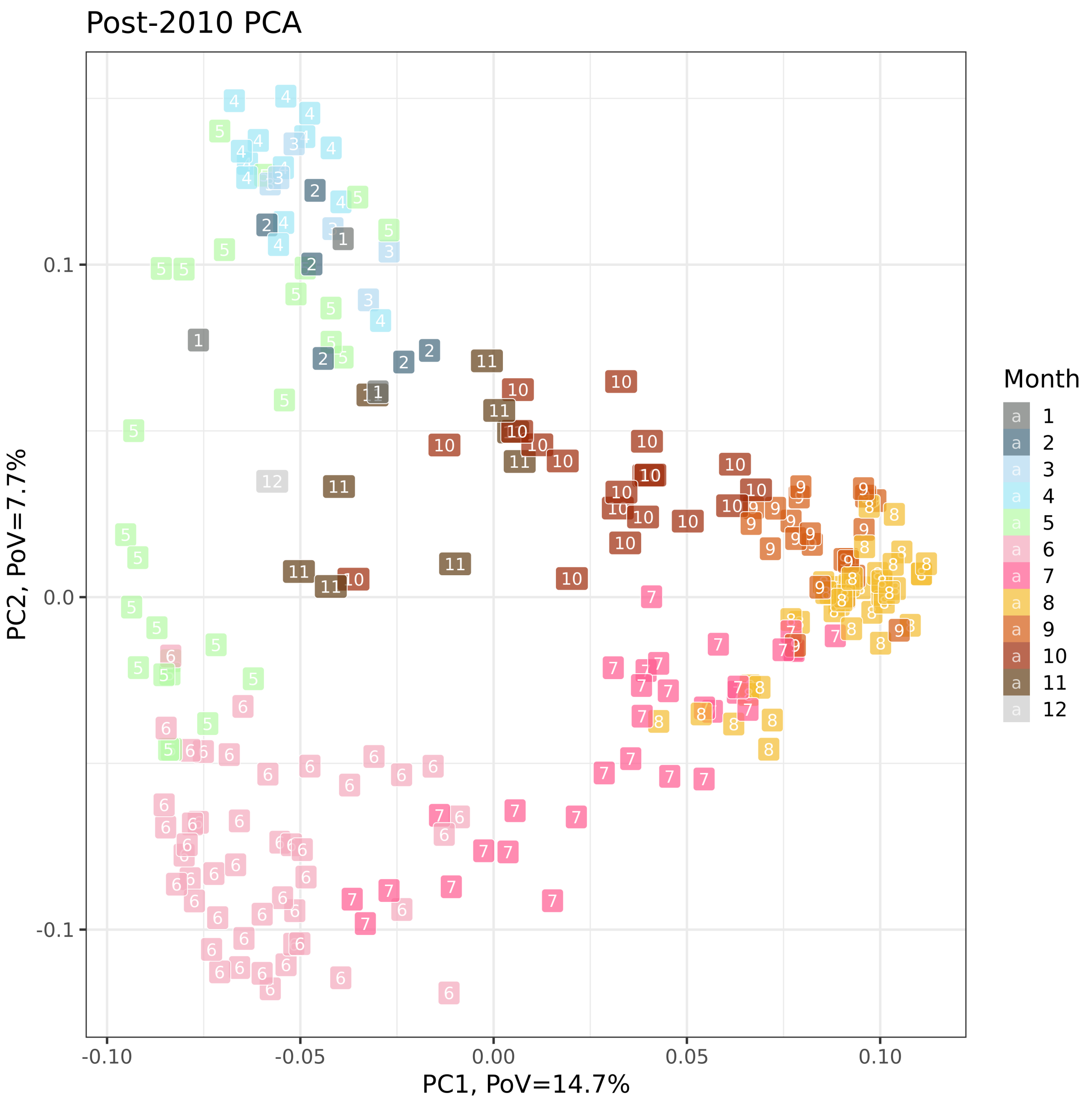


Figure 9: PCA on datapoints after 2010 showing partitioning of samples according to season despite the fact that the PCA showed a low proportion of variance explained for the first two principal components (shown on axes; principal component 1 explained 14.7% of the variability in community abundance profiles while principal component 2 explained 7.7% of the variability). This is in line with the complexity of the data, making it challenging to assign only two axes of major variability.

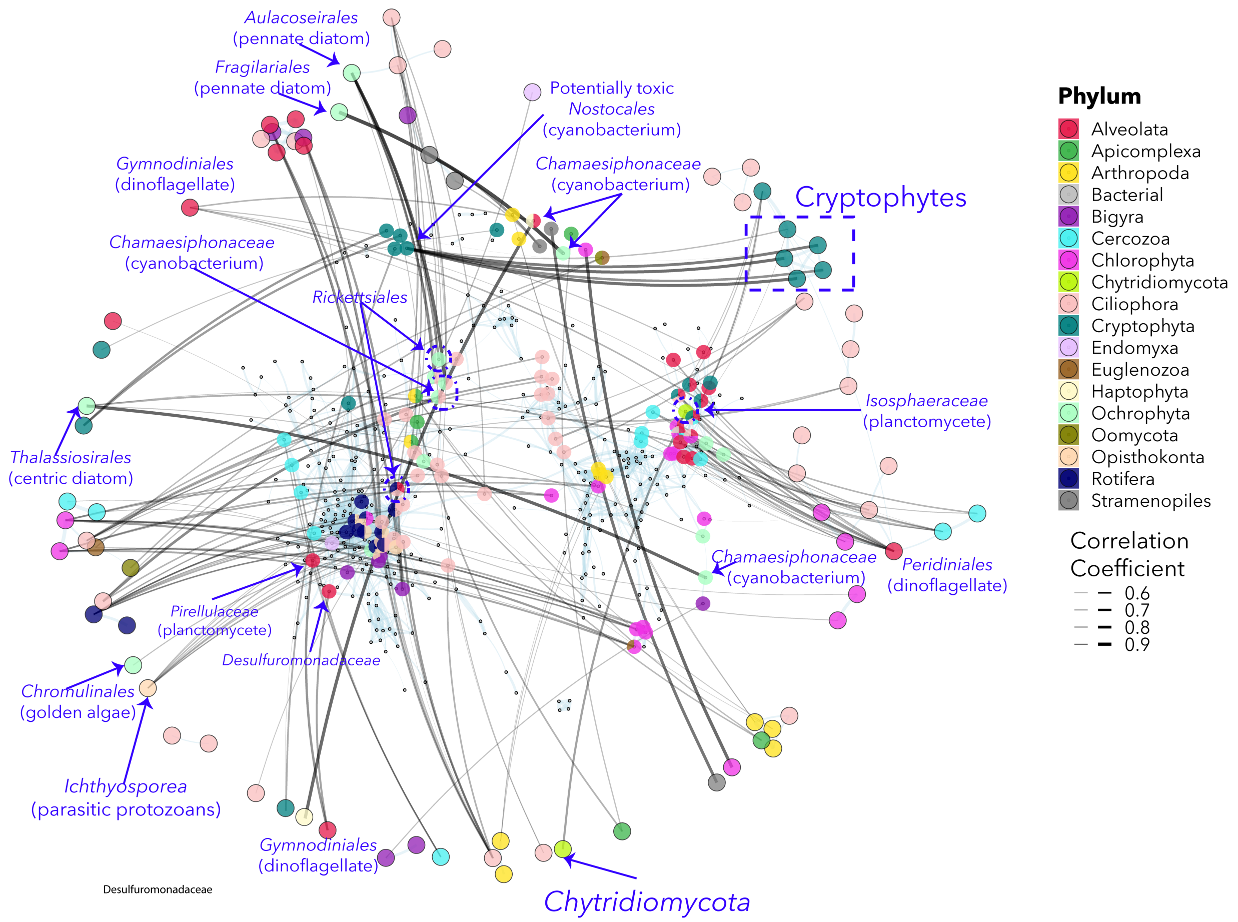


Figure 10: **Network analysis over 20-year time series of metagenomes suggests stable connections between some eukaryotes and bacteria.** Only correlations with a significant Benjamini-Hochberg-corrected p-value and a correlation coefficient of greater than or equal to 0.50 are included visually. A selection of interactions are labeled on the network. The outer circle of the network is a correlation-weighted representation of eukaryote-eukaryote interactions, while the inner circle of the network is a representation of bacterial relationships. All within-domain correlations are in light blue. The black lines have thickness corresponding to the correlation coefficient of bacteria-eukaryote correlations. Bacteria that have correlations to eukaryotes have colored circles over top of the underlying point in the bacteria-bacteria network; each of these circles is a pie chart colored according to the taxonomic phylum of the eukaryote(s) that they correlate to.
